# Integration of machine learning and genome-scale metabolic modeling identifies multi-omics biomarkers for radiation resistance

**DOI:** 10.1101/2020.08.02.233098

**Authors:** Joshua E. Lewis, Melissa L. Kemp

## Abstract

Resistance to ionizing radiation, a first-line therapy for many cancers, is a major clinical challenge. Personalized prediction of tumor radiosensitivity is not currently implemented clinically due to insufficient accuracy of existing machine learning classifiers. Despite the acknowledged role of tumor metabolism in radiation response, metabolomics data is rarely collected in large multi-omics initiatives such as The Cancer Genome Atlas (TCGA) and consequently omitted from algorithm development. In this study, we circumvent the paucity of personalized metabolomics information by characterizing 915 TCGA patient tumors with genome-scale metabolic Flux Balance Analysis models generated from transcriptomic and genomic datasets. Novel metabolic biomarkers differentiating radiation-sensitive and -resistant tumors were predicted and experimentally validated, enabling integration of metabolic features with other multi-omics datasets into ensemble-based machine learning classifiers for radiation response. These multi-omics classifiers showed improved classification accuracy, identified novel clinical patient subgroups, and demonstrated the utility of personalized blood-based metabolic biomarkers for radiation sensitivity. The integration of machine learning with genome-scale metabolic modeling represents a significant methodological advancement for identifying prognostic metabolite biomarkers and predicting radiosensitivity for individual patients.

## Introduction

Despite being one of the oldest forms of cancer therapy and still a primary treatment modality, radiation therapy is not effective for over one-fifth of cancer patients distributed across almost all cancer types^1, 2^. While biological understanding of radiation resistance has been advanced, use of *a priori* prediction of radiation response for individual cancer patients is not yet implemented clinically^3^. Early studies that identified biomarkers for radiation response focused on tumor histology, clinical factors including staging and Karnofsky performance score, and physiological parameters such as tumor oxygenation status^4–6^. As methods for transcriptomic analysis have improved, gene expression-based classifiers for radiation response have proliferated (recently curated in the RadiationGeneSigDB database)^7^. To date, however, these radiation response classifiers do not integrate multiple –omics modalities, owing in part to a lack of available -omics datasets for individual patient tumors. Specifically, while genomic and transcriptomic data is becoming more widely available for large numbers of patient tumors through initiatives such as The Cancer Genome Atlas (TCGA), metabolomic data associated with tumor biobanks is rarely captured, limiting inclusion of tumor metabolic features in predictive models for radiation therapy response^2^.

Given the lack of available tumor metabolomic data, genome-scale metabolic modeling approaches such as flux balance analysis (FBA) are becoming increasingly popular for predicting metabolic phenotypes^8, 9^. By combining a curated reconstruction of the human metabolic network with constraints on metabolic reaction activities and an objective function to maximize a particular metabolic phenotype, predictions of steady-state reaction fluxes or metabolite production rates under physiological constraints can be obtained at a genome scale^10^. We previously developed a novel bioinformatics pipeline for integrating genomic, transcriptomic, kinetic, and thermodynamic parameters into personalized FBA models of 716 radiation-sensitive and 199 radiation-resistant patient tumors from TCGA across multiple cancer types^11^. Using these metabolic models, we identified novel differences in redox metabolism between radiation-sensitive and -resistant tumors, as well as personalized gene targets for inhibiting antioxidant production and clearance of reactive oxygen species. By validating model predictions using a panel of matched radiation-sensitive and -resistant cancer cell lines, we demonstrated that genome-scale metabolic models provide accurate predictions of tumor metabolism and can identify diagnostic and therapeutic biomarkers for radiation response.

While machine learning methods have been previously combined with genome-scale metabolic models to improve prediction of metabolic phenotypes, most studies combining these two methodologies have focused on microbiological applications rather than applications to cancer metabolism or predicting treatment outcomes^12^. We hypothesize that predictions from genome-scale metabolic models of patient tumors would provide additional information for distinguishing pathophysiological differences between radiation-sensitive and -resistant tumors, as well as for prediction of radiation of radiation response. To this end, we utilized personalized FBA models of TCGA patient tumors to predict genome-scale metabolite production rates for incorporation into machine learning classifiers and identifying novel metabolite biomarkers associated with radiation resistance. Additionally, through integration with clinical, genomic, and transcriptomic datasets, we developed gene expression, multi-omics, and non-invasive classifiers which outperform previous predictors of radiation response, as well as provide personalized diagnostic biomarker panels for individual patient tumors.

## Results

### Gene expression classifier implicates cellular metabolism

Because the majority of previously-developed classifiers for radiation response are based on gene expression data (curated in the RadiationGeneSigDB database), we first developed a machine learning classifier utilizing transcriptomic data from radiation-sensitive and -resistant TCGA tumors to compare predictive accuracy and identified gene sets^7^. A gradient boosting machine (GBM) algorithm was used with Bayesian optimization for determining optimal hyperparameter values, providing optimal performance accuracy on TCGA datasets (**Supplementary Fig. 1**). 782 of the 22,819 genes in the dataset (3.43%) were identified as significant in the classification of radiation response, determined by a 95% cumulative sum threshold on feature importance scores (**Fig. 1a**). 10 of the 50 genes with largest feature importance scores were previously implicated in radiation therapy response^13–22^. To determine whether the identified set of 782 genes has more predictive value than previously-identified gene sets in RadiationGeneSigDB, machine learning classifiers were subsequently trained using only the genes from each respective gene set, and the predictive accuracy of each classifier was compared (**Fig. 1b**). Our set of 782 genes had the best performance among all gene sets when trained on the TCGA dataset, and was among the best gene sets when trained on a separate dataset from the Cancer Cell Line Encyclopedia (CCLE)^23^.

**Fig. 1.**
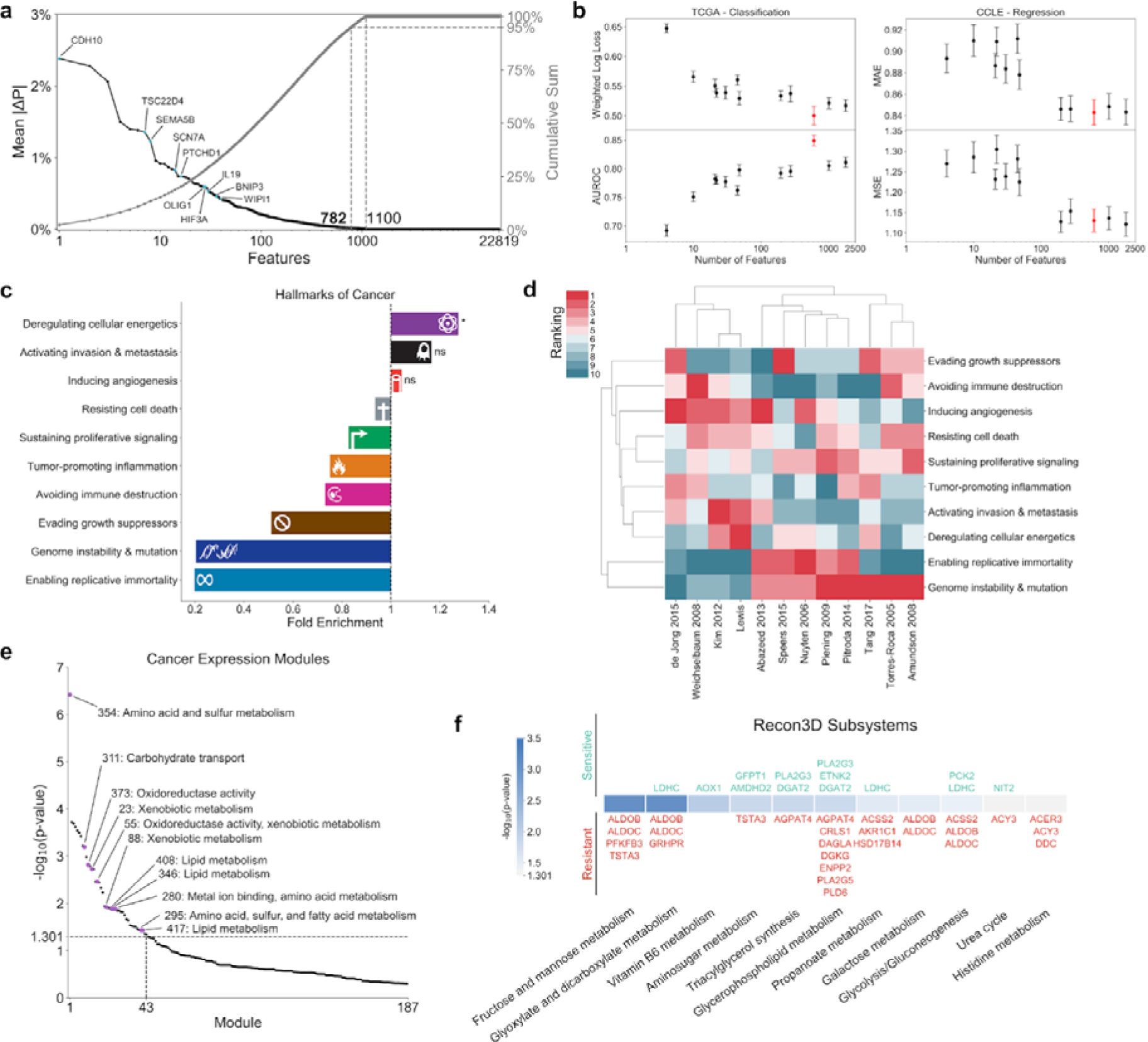
Gene expression classifier for radiation response. **a**, (Left, black) Feature importance scores for individual genes, signifying the absolute change in predicted probability of radiation resistance attributed to each feature averaged across all samples. Those features within the top 50 with previous literature suggesting a role in tumor radiation response are annotated. (Right, gray) Cumulative feature importance scores. **b,** Performance of the identified set of 782 significant gene expression features from this study (red) versus previously identified gene sets in RadiationGeneSigDB (black), on both the (left) TCGA dataset performing a classification task on patient tumor radiation response and (right) CCLE dataset performing a regression task on cancer cell line radiation response. AUROC: area under the receiver operating characteristic curve; MAE: mean absolute error; MSE: mean squared error. **c,** Gene set enrichment analysis (GSEA) of significant features from our gene expression classifier among the Hallmarks of Cancer. **d,** Hierarchical clustering of Hallmarks of Cancer enrichment ranks from the gene set in this study and those in RadiationGeneSigDB, based on both (row) hallmark, and (column) gene set. **e,** GSEA of significant gene expression features among the cancer expression modules from Segal et al. Modules relevant to cellular metabolism are annotated with their number and descriptions. **f,** GSEA of significant gene expression features among Recon3D metabolic subsystems. Significant genes within each subsystem are annotated above or below p-value bars based on whether their expression is positively correlated with (above, green) radiation sensitivity, or (below, red) radiation resistance. ns: not significant, *: p ≤ 0.05, **: p ≤ 0.01, ***: p ≤ 0.001, ****: p ≤ 0.0001.

Gene set enrichment analysis (GSEA) of these 782 genes among the Hallmarks of Cancer showed significantly increased enrichment of the “Deregulating cellular energetics” hallmark, with very low enrichment of the “Genome instability & mutation” hallmark (**Fig. 1c**)^24, 25^. Hierarchical clustering of the hallmark enrichment ranks for each gene set in RadiationGeneSigDB revealed two major clusters: a larger cluster with very high rank of “Genome instability & mutation”, and a smaller cluster with much higher ranks for other hallmarks involved in cellular metabolism, angiogenesis, and metastasis (**Fig. 1d**). This dichotomy suggests that although the biological response to radiation therapy certainly involves genomic instability and DNA-damage repair, other biological processes such as cellular metabolism may play critical roles as well^26, 27^. GSEA of cancer expression modules additionally showed increased enrichment of many modules involved in cellular metabolism, including amino acid and sulfur metabolism, redox metabolism, and lipid metabolism (**Fig. 1e**)^28^. Finally, GSEA of Recon3D metabolic subsystems demonstrated increased enrichment of pathways involved in central carbon metabolism and lipid metabolism, with the majority of genes being associated with increased probability of radiation resistance (**Fig. 1f**)^10^. Together, analysis of this gene expression classifier suggests that radiation-resistant tumors exemplify dysregulation in their cellular metabolic networks, and that additional features involving the metabolism of radiation-sensitive and -resistant tumors will provide significant benefit in developing machine learning classifiers for radiation response.

### FBA models accurately predict relative metabolite production

Personalized genome-scale FBA models of radiation-sensitive and -resistant TCGA tumors were generated to obtain metabolic features which could be used in machine learning classifiers for radiation response. These FBA models were developed through integration of gene expression and mutation information from individual patient tumors, as well as kinetic and thermodynamic parameters frompublicly-available repositories^11^. By systematically creating artificial metabolite sinks in the Recon3D metabolic network and evaluating fluxes to these sinks, the production rates of different metabolites were predicted and compared between radiation-sensitive and -resistant tumors (**Fig. 2a**). **Fig. 2b** shows that many of the metabolite classes implicated from the gene expression classifier showed significantly increased production in radiation-resistant tumors. These included antioxidant and cysteine-containing metabolites (including precursors of glutathione, an antioxidant with previously-implicated roles in radiation response)^29^, lipid and fatty acid metabolites (including those previously implicated in lipid peroxidation in response to ionizing radiation)^30, 31^, and immune system mediators. While fewer metabolites were predicted to be significantly downregulated in radiation-resistant tumors, many metabolites involved in nucleotide metabolism were among this group.

**Fig. 2.**
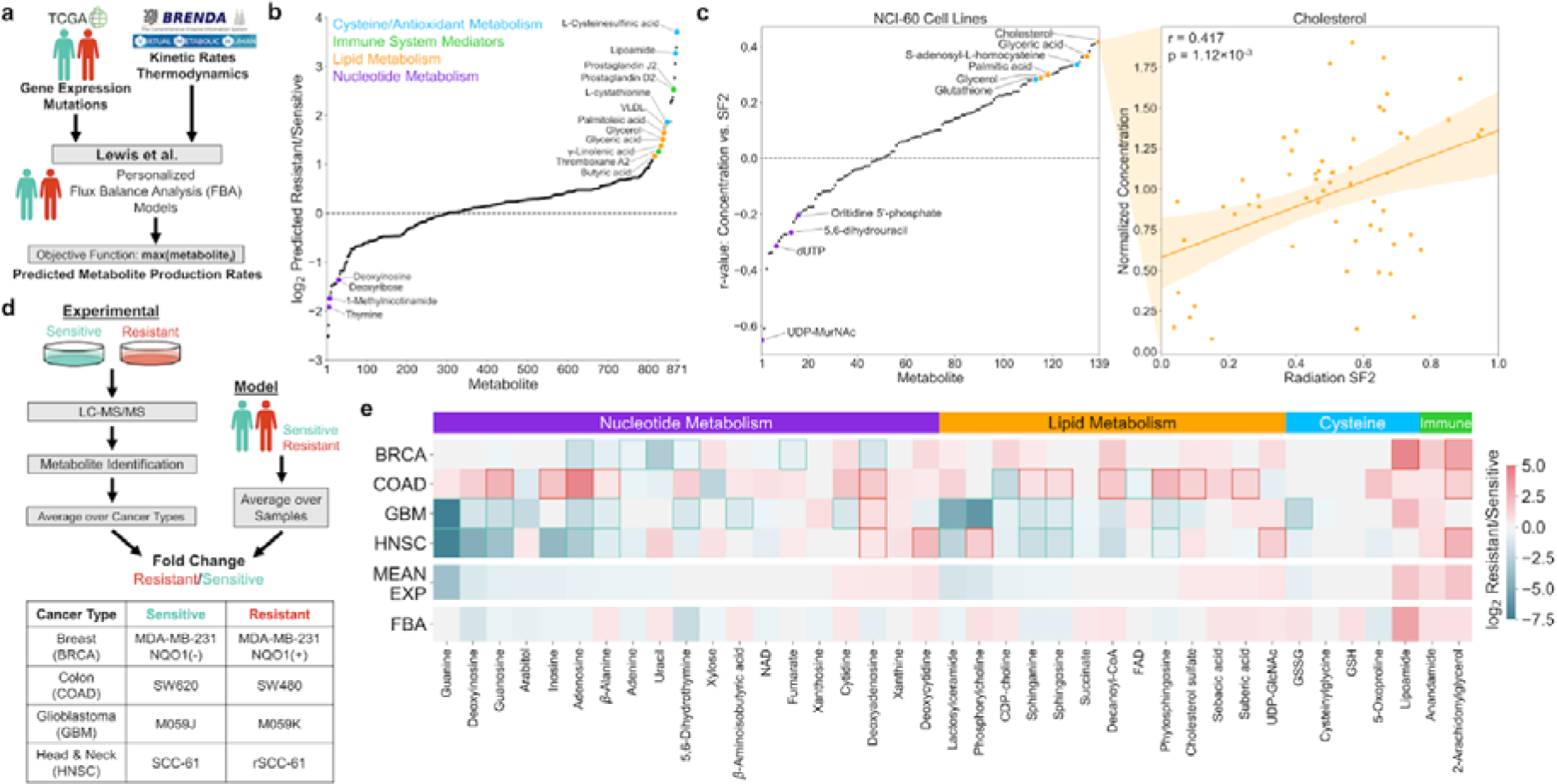
FBA model predictions of relative metabolite production and experimental validation between radiation-sensitive and radiation-resistant cancers. **a**, Multi-omics data from TCGA tumors and publicly-available repositories is integrated to develop personalized FBA models and predict differences in metabolite production rates between radiation-sensitive and -resistant tumors. **b,** Model-predicted metabolite production rates, expressed as the log_2_ ratio of average production between radiation-resistant versus -sensitive tumors. Metabolites within major classes with significant upregulation or downregulation in radiation-resistant tumors are color-coded and annotated. **c,** (Left) Correlation between metabolite concentration and surviving fraction at 2 Gy radiation (SF2) among 139 experimentally-measured metabolites in the NCI-60 panel of cancer cell lines. Metabolite classes are colored as in (**b**). (Right) Example regression between metabolite concentration and cell line SF2 for cholesterol. **d,** (Top) Schematic showing the comparison of model-predicted metabolite production in radiation-sensitive and -resistant TCGA tumors, with experimentally-measured metabolite concentrations in matched radiation-sensitive and -resistant cell lines. (Bottom) Radiation-sensitive and -resistant cell line pairs across four different cancer types used in the experimental metabolomics study (**Supplementary Table 1**). **e,** Comparison of model-predicted and experimentally-measured levels of individual putative metabolites within the four major classes identified in (**b**). BRCA, COAD, GBM, HNSC: log_2_ ratio of putative metabolite levels in radiation-resistant versus -sensitive cell lines. Statistically-significant differences within each cell line pair are represented by box outlines. MEAN EXP: average experimental log_2_ ratio across all four cell line pairs. FBA: log_2_ ratio of model-predicted metabolite production rates in radiation-resistant versus -sensitive TCGA tumors.

Regression of experimental metabolite concentrations among the NCI-60 cancer cell line panel with cell line surviving fraction at 2 Gy radiation (SF2) showed up- and down-regulation of the same metabolite classes predicted from FBA models (**Fig. 2c**)^32^. Many lipid and fatty acid metabolites positively correlate with radiation resistance (including cholesterol, which had the most positive correlation among all metabolites tested); antioxidant metabolites including glutathione positively correlate as well. On the other hand, many nucleotide metabolites are anti-correlated with radiation resistance (including UDP-MurNAc, which had the most negative correlation among all metabolites tested).

To experimentally validate FBA model predictions of individual metabolite levels, we analyzed matched pairs of radiation-sensitive and -resistant cell lines from four different cancer types via untargeted metabolomics (**Fig. 2d-e, Supplementary Fig. 2-5, Supplementary Table 1**). The pan-cancer FBA models accurately predicted that most nucleotide metabolites, including derivatives of adenine, guanine, thymine, and inosine, are downregulated in radiation-resistant cancers, while, in contrast, cytidine derivatives are upregulated. Predictions of lipid production accurately captured the observed heterogeneity in lipid levels between cell lines. Although model-predictions of absolute oxidized (GSSG) and reduced (GSH) glutathione production did not match with experimentally-measured values, previous model predictions of increased reduction potential of GSSG to GSH in radiation-resistant tumors agreed with experimental findings of greater GSH/GSSG ratios in radiation-resistant cell lines^11^. Finally, model-predicted production of the antioxidant lipoamide as well as immune mediators anandamide and 2-arachidonylglycerol corresponded very well with experimental measurements, which were upregulated in nearly all radiation-resistant cell lines. Overall, these findings demonstrate that genome-scale metabolic models derived from transcriptomic and genomic data provide surprisingly accurate predictions of relative metabolite production between radiation-sensitive and -resistant cancers, allowing for their use in machine learning classifiers for radiation response.

### Machine learning architecture for radiation response

To integrate FBA model predictions of metabolite production rates with other TCGA datasets into multi-omics machine learning classifiers, a dataset-independent ensemble architecture was developed (**Fig. 3a**). Multiple independent “base learner” classifiers are trained on an individual -omics dataset (either clinical, genomics, transcriptomics, or metabolomics data), as described in **Supplementary Fig. 1**. Subsequently, by comparing predicted class probabilities from each individual base learner to known radiation responses, a “meta-learner” classifier is trained to determine which base learner provides the most accurate prediction of radiation response based on the multi-omics features of individual samples (**Fig. 3b**)^33^. For an individual testing sample, each base learner outputs the predicted probability of radiation resistance (*p_i_*), and the meta-learner outputs the predicted probability that each base learner will provide the most accurate prediction (*w_i_*); the final probability of radiation resistance is the weighted average of each *p_i_*, with weights being each *w_i_* (**Fig. 3c**). This dataset-independent ensemble architecture performs better across multiple performance metrics compared to the common practice of initially combining all -omics datasets and training on a single classifier (**Fig. 3d, Supplementary Fig. 6-7**). Overall, this machine learning architecture is a robust platform for integrating multi-omics data and providing accurate predictions of radiation response in individual patient tumors.

**Fig. 3.**
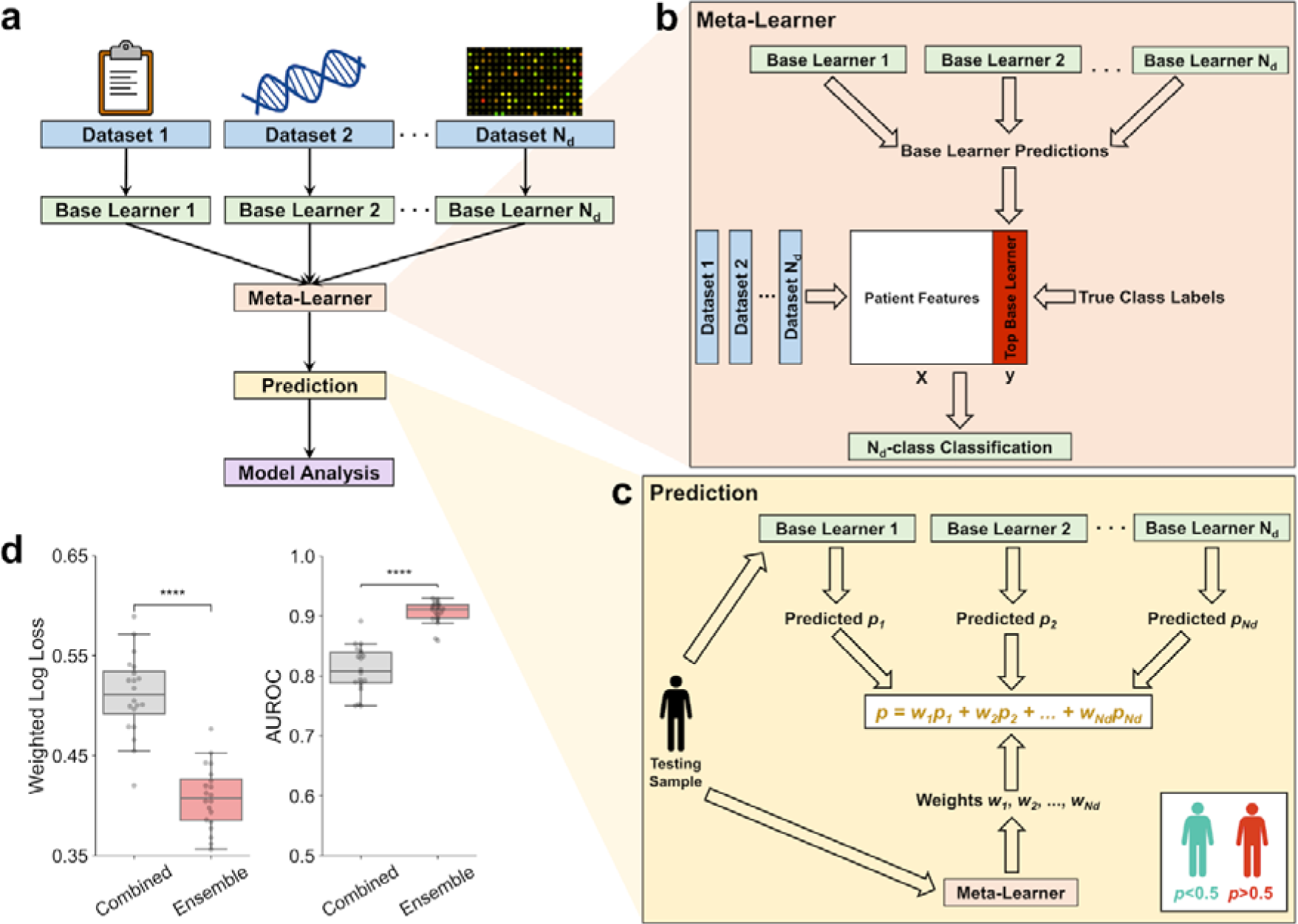
Machine learning architecture for improved prediction of radiation therapy response. **a**, Dataset-independent ensemble architecture, with independent base learners for each dataset and one meta-learner for integration of base learner outputs. **b,** Meta-learner performing N_d_-class classification of the most accurate base learner/dataset for each sample, where N_d_ is the number of independent base learners/datasets. **c,** Prediction of radiation response for each testing set sample using predicted probabilities from each base learner and weights from the meta-learner. **d,** Performance of multi-omics classifier trained on clinical, gene expression, mutation, and FBA-predicted metabolite data from TCGA tumors, comparing the dataset-independent ensemble architecture versus combining datasets together before training on a single classifier. Weighted log loss and AUROC performance metrics are shown here, with other metrics shown in **Supplementary Fig. 6**. ns: not significant, *: p≤ 0.05, **: p≤0.01, ***: p 0.001, ****: p≤ ≤ 0.0001.

### Multi-omics classifier identifies clinical patient subgroups

Using the dataset-independent ensemble architecture described above, a multi-omics machine learning classifier integrating clinical, gene expression, mutation, and FBA-predicted metabolite production rates from TCGA tumors was developed. With an AUROC of 0.906 ± 0.004, this classifier has significantly greater performance compared to previously-developed machine learning classifiers for radiation response (**Fig. 4a, 1b**)^7, 34^. Additionally, the threshold for separating radiation-sensitive and -resistant classes can be altered to optimize sensitivity, specificity, or a balance of both. 725 of the 52,223 features from the four datasets (1.39%) were identified as significant in the classification of radiation response (**Fig. 4b**). While the majority of these 725 features were gene expression (48.3%) and metabolite (32.6%) features, clinical features including tumor histology, chemotherapeutic response, and cancer type contributed more than half of the total feature importance scores (60.1%; **Fig. 4c**). Mutations with significant feature importance scores included those directly involved in redox metabolism (*IDH1* R132H) and lipid metabolism (*BRAF* V600E)^35, 36^.

**Fig. 4.**
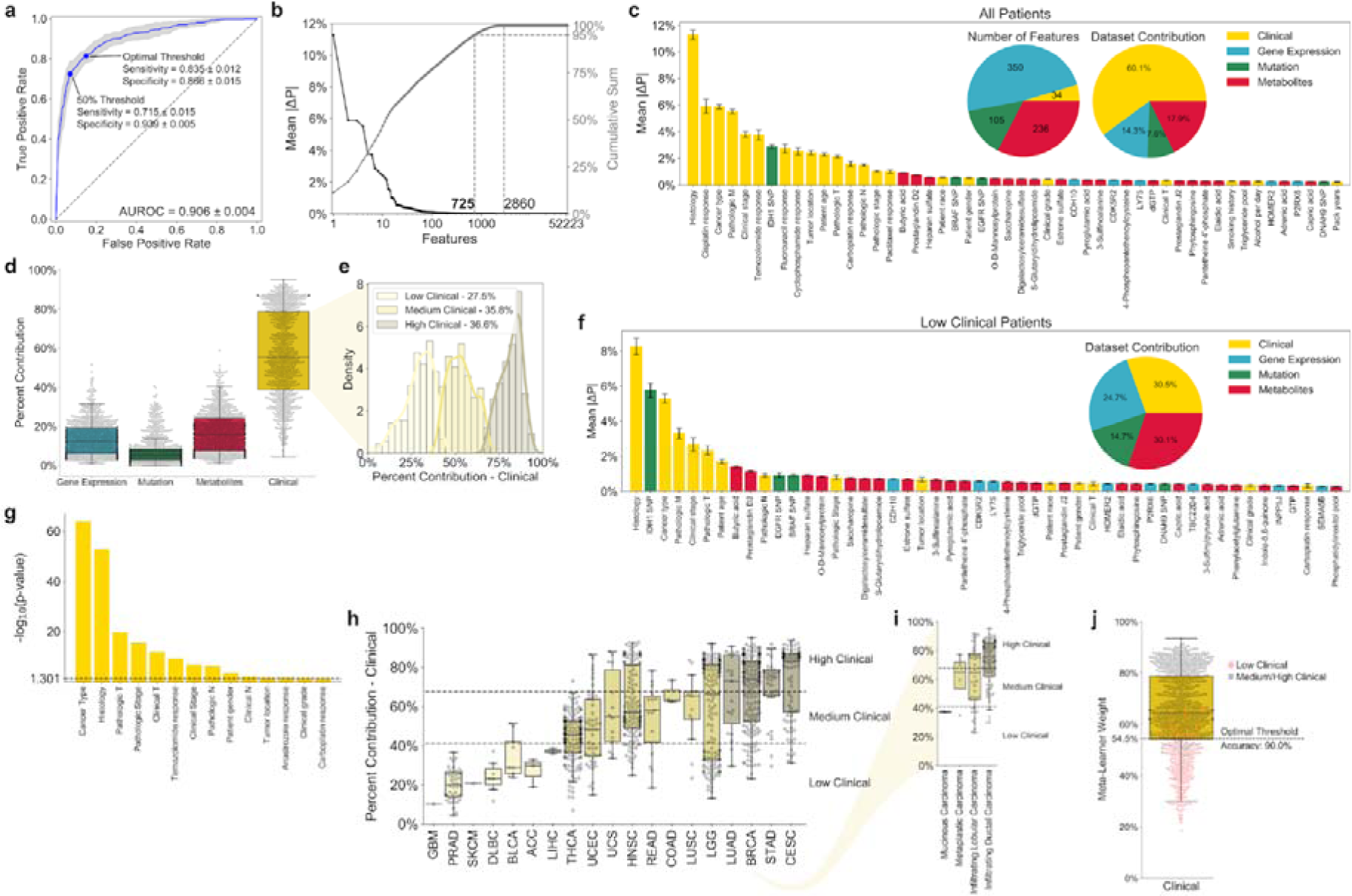
Multi-omics classifier integrating clinical, gene expression, mutation, and FBA-predicted metabolite features for prediction of radiation response. **a**, Receiver operating characteristic (ROC) curve for the multi-omics classifier. The point representing a 50% predicted probability threshold for separating radiation-sensitive and -resistant classes, as well as the optimal point for maximizing Youden’s J statistic (J = sensitivity + specificity - 1) are shown. Blue line: mean across 20 training+validation/testing splits. Light blue error band: ± 1 standard deviation. **b,** (Left, black) Feature importance scores for individual features. (Right, gray) Cumulative feature importance scores. **c,** List of top 50 features with largest feature importance scores, colored based on their original dataset. (Inset, Left) Number of significant features from each dataset. (Inset, Right) Relative contribution of features from each dataset to the sum of total feature importance scores, averaged across all samples. **d,** Relative contribution of features from each dataset to the sum of total feature importance scores, for each individual sample. **e,** Clustering of samples into “Low”, “Medium”, and “High” clinical groups based on the relative contribution of the clinical dataset. The optimal number of clusters was calculated based on maximizing the gap statistic from k-means clustering (**Supplementary Fig. 8a**). **f,** Top 50 features with largest feature importance scores among samples within the “Low Clinical” cluster. (Inset) Relative contribution of features from each dataset to the sum of total feature importance scores, averaged across all samples within the “Low Clinical” cluster. **g,** Statistical significance of patient clustering into “Low/Medium/High” clinical groups based on different clinical factors, as calculated by the chi-squared test with Yates’ correction. Only statistically significant (p ≤ 0.05) clinical factors are shown. **h,** Clinical cluster and dataset contribution of samples within different cancer types. **i,** Clinical cluster and dataset contribution of breast cancer (BRCA) samples with different histological subtypes. **j,** Prediction of clinical cluster based on meta-learner weight for the clinical dataset. Dotted line: threshold which maximizes the accuracy in separating “Low Clinical” from “Medium/High Clinical” clusters.

Individual samples varied significantly in the contribution of different datasets towards radiation response classification (**Fig. 4d**). Using unsupervised clustering, three clusters of patients with varying contributions of clinical features were identified (**Fig. 4e, Supplementary Fig. 8a**). While “High Clinical” patients were categorized by large clinical feature contributions and small contributions from multi-omics datasets, multi-omics features provided the majority of cumulative feature importance scores for “Low Clinical” patients, with metabolic features alone providing nearly as much utility as clinical features (**Fig. 4f**). For this “Low Clinical” cluster, certain clinical features including chemotherapeutic response have diminutive utility, whereas novel multi-omics features including *IDH1* SNP and lipid metabolite levels have much higher importance scores compared to the overall patient cohort. Significant heterogeneity in clinical clusters was observed based on patient clinical factors, especially cancer type and tumor histology (**Fig. 4g-i**). Output weights from the meta-learner provide an accurate prediction of clinical cluster, effectively differentiating between “Low Clinical” and “Medium/High Clinical” patients; this provides a valuable strategy for determining whether clinical information from electronic medical records is sufficient to accurately predict radiation response in an individual patient, or whether multi-omics features from tumor biopsy samples are needed (**Fig. 4j**).

### Novel metabolic biomarkers identified for radiation response

Metabolite set enrichment analysis (MSEA) of the 236 significant metabolite features from the multi-omics classifier indicated significant enrichment of several metabolic pathways involved in central carbon metabolism, lipid metabolism, and nucleotide metabolism (**Fig. 5a**). To identify individual metabolites with the largest impact on radiation response prediction, the Spearman correlation between feature importance score and predicted metabolite production rate across all patients was calculated for each metabolite (**Supplementary Fig. 9**). **Fig. 5b** highlights many of the significant metabolic features, as well as metabolism-related gene expression and mutation features. Significant glycolytic and TCA cycle metabolites (fructose 1,6-bisphosphate, 3-phosphoglyceric acid, succinyl-CoA, and succinate) were all positively correlated with radiation resistance, while genes promoting gluconeogenesis (*PCK2* and *LDHC*) were associated with radiation sensitivity. Fructose 2,6-bisphosphate, an allosteric regulator of PFK-1 that activates glucose breakdown, had one of the most positive correlation values. Additionally, many metabolites in early mannose metabolism had positive correlation values, in accordance with previously observed radiation-induced upregulation of mannose-6-phosphate receptors and high-mannose type N-glycan production^37, 38^.

**Fig. 5.**
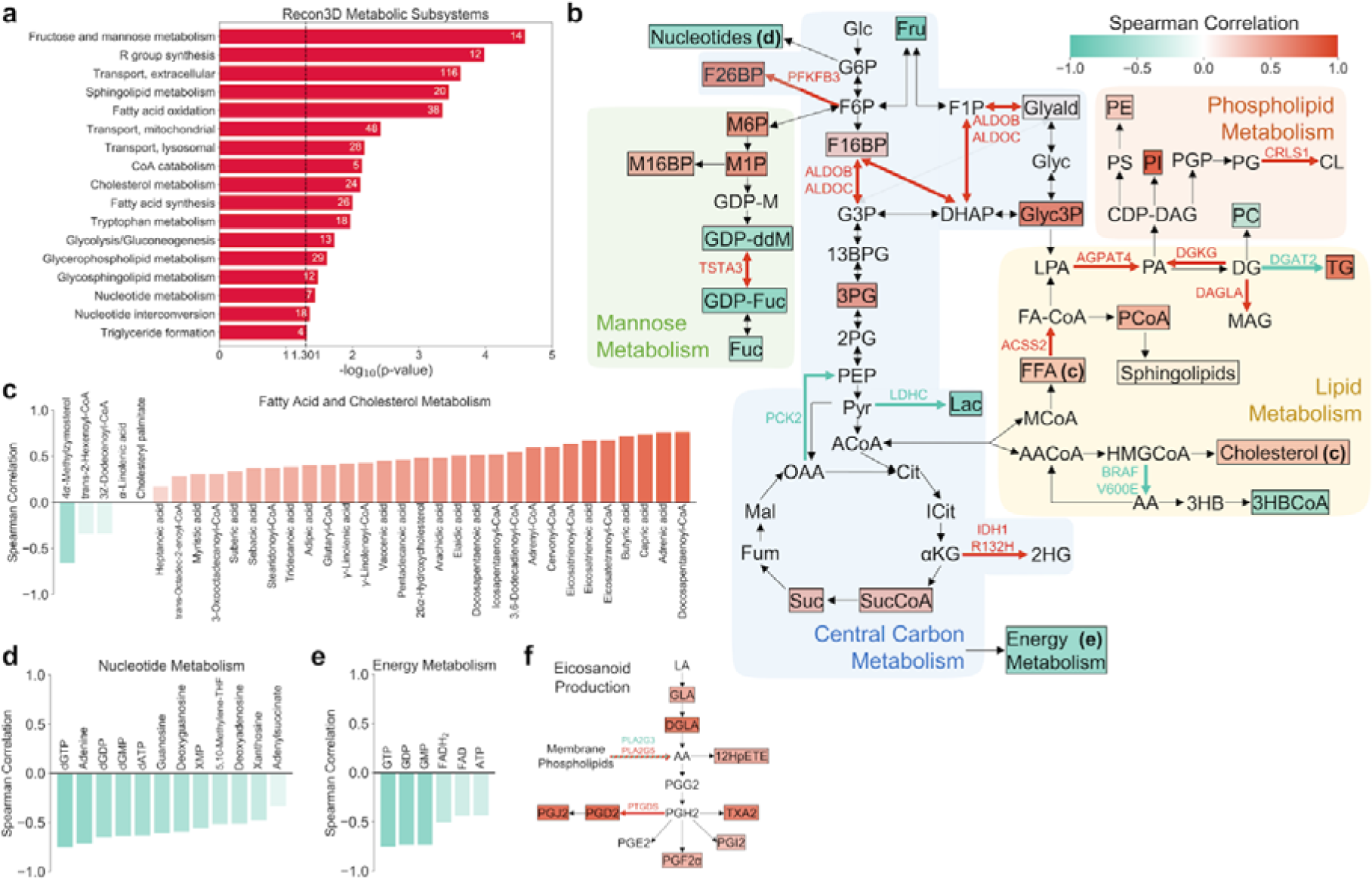
Analysis of metabolic biomarkers from the multi-omics classifier for radiation response. **a**, Metabolite-set enrichment analysis (MSEA) of significant metabolic features among Recon3D metabolic subsystems. The number of significant metabolites in each subsystem are shown. Only statistically significant (p ≤ 0.05) subsystems are shown. b, Overview of significant metabolic features, as well ≤ as metabolism-related gene expression and mutation features. Different metabolic pathways are shown with colored backgrounds. Significant metabolic features are denoted by colored boxes, where the color indicates the Spearman correlation coefficient between feature importance score and predicted metabolite production rate across all patients (**Supplementary Fig. 9**). Significant gene expression and mutation features are denoted by colored reaction arrows, either in green (associated with radiation sensitivity) or in red (associated with radiation resistance). 13BPG: 1,3-bisphosphoglycerate; 2HG: 2-hydroxyglutarate; 2PG: 2-phosphoglycerate; 3HB: 3-hydroxybutyrate; 3HBCoA: 3-hydroxybutyrl-CoA; 3PG: 3-phosphoglycerate; KG: Alpha-ketoglutarate; AA: Acetoacetate; AACoA: αAcetoacetyl-CoA; ACoA: Acetyl-CoA; CDP-DAG: CDP-diacylglycerol; Cit: Citrate; CL: Cardiolipin; DG: Diacylglycerol; DHAP: Dihydroxyacetone phosphate; F16BP: Fructose 1,6-bisphosphate; F1P: Fructose 1-phosphate; F26BP: Fructose 2,6-bisphosphate; F6P: Fructose 6-phosphate; FA-CoA: Fatty acyl-CoA; FFA: Free fatty acid; Fru: Fructose; Fuc: Fucose; Fum: Fumarate; G3P: Glyceraldehyde 3-phosphate; G6P: Glucose 6-phosphate; GDP-ddM: GDP-4-keto-6-deoxymannose; GDP-Fuc: GDP-fucose; GDP-M: GDP-mannose; Glc: Glucose; Glyald: Glyceraldehyde; Glyc3P: Glycerol 3-phosphate; Gylc: Glycerol; HMGCoA: 3-hydroxy-3-methylglutaryl-CoA; ICit: Isocitrate; Lac: Lactate; LPA: Lysophosphatidic acid; M16BP: Mannose 1,6-bisphosphate; M1P: Mannose 1-phosphate; M6P: Mannose 6-phosphate; MAG: Monoacylglycerol; Mal: Malate; MCoA: Malonyl-CoA; OAA: Oxaloacetate; PA: Phosphatidic acid; PC: Phosphatidylcholine; PCoA: Palmitoyl-CoA; PE: Phosphatidylethanolamine; PEP: Phosphoenolpyruvate; PG: Phosphatidylglycerol; PGP: Phosphatidylglycerol-phosphate; PI: Phosphatidylinositol; PS: Phosphatidylserine; Pyr: Pyruvate; Suc: Succinate; SucCoA: Succinyl-CoA; TG: Triglyceride. **c-e,** Spearman correlation coefficients of significant metabolic features involved in (**c**) fatty acid and cholesterol metabolism, (**d**) nucleotide metabolism, and (**e**) energy metabolism. **f,** Metabolic pathway of eicosanoid production, highlighting significant metabolite and gene expression features. 12HpETE: 12-hydroxyperoxyeicosatetraenoic acid; AA: Arachidonic acid; DGLA: Dihomo-γ-linolenic acid; GLA: γ-linolenic acid; LA: linoleic acid.

Greater glycolytic fluxes in radiation-resistant tumor models resulted in increased production of the majority of significant lipid and fatty acid metabolites, including many with previously-identified roles in antioxidation such as capric acid, butyric acid, eicosatrienoic acid, and γ-linolenic acid (**Fig. 5c**) ^39–42^. On the other hand, significant nucleotide metabolites were highly correlated with radiation sensitivity, in agreement with the observed downregulation in radiation-resistant cancer cell lines (**Fig. 5d**). While production of energy metabolites including ATP was correlated with radiation sensitivity, FBA models predict significantly greater conversion of ADP to ATP in radiation-resistant tumors, in agreement with previous experimental findings (**Fig. 5e, Supplementary Fig. 10**)^43, 44^. Finally, increased production of membrane phospholipids and arachidonic acid precursors resulted in significant correlations between inflammation-mediating eicosanoids and radiation resistance, corroborating previous evidence of radiation-sensitizing effects of cyclooxygenase inhibitors including aspirin (**Fig. 5f**)^45^. Together, these findings suggest that metabolic features from multiple interconnected pathways including central carbon, lipid, and nucleotide metabolism are viable diagnostic biomarkers for prediction of radiation sensitivity.

### Non-invasive classifier implicates blood metabolic features

Because non-invasive metabolic predictors of radiation response could be rapidly applied for informing patient-specific treatment, we refined machine learning classification to only integrate clinical data derived from non-invasive means (excluding any pathologic staging or histological information from tumor biopsies) with FBA-predicted production rates of metabolites known to be quantifiable in human blood samples (**Fig. 6a**)^46^. This non-invasive classifier performed similarly overall to the multi-omics classifier, with increased sensitivity and decreased specificity; this suggests that the non-invasive classifier may be optimal as a first-line screening test, followed by the multi-omics classifier as a second-line diagnostic test (**Fig. 6b**)^47^. 97 of the 363 features from the two datasets (26.7%) were identified as significant in the classification of radiation response (**Fig. 6c**). Similar to the multi-omics classifier (**Fig. 4e**), individual patient contributions of clinical features formed a bimodal distribution of “Low Clinical” and “High Clinical” groups (**Fig. 6d, Supplementary Fig. 8b**). Blood metabolite features - including many lipid, nucleotide, and inflammation-mediating metabolites previously identified from the multi-omics classifier - provided almost one-half of the cumulative feature importance scores for “Low Clinical” patients (**Fig. 6e**). Dataset contributions and feature importance scores for individual cancer patients can identify personalized biomarkers with maximal diagnostic utility (**Fig. 6f-h**). Overall, these findings demonstrate the value of blood-based biomarkers as a non-invasive approach towards personalized prediction of radiation response.

**Fig. 6.**
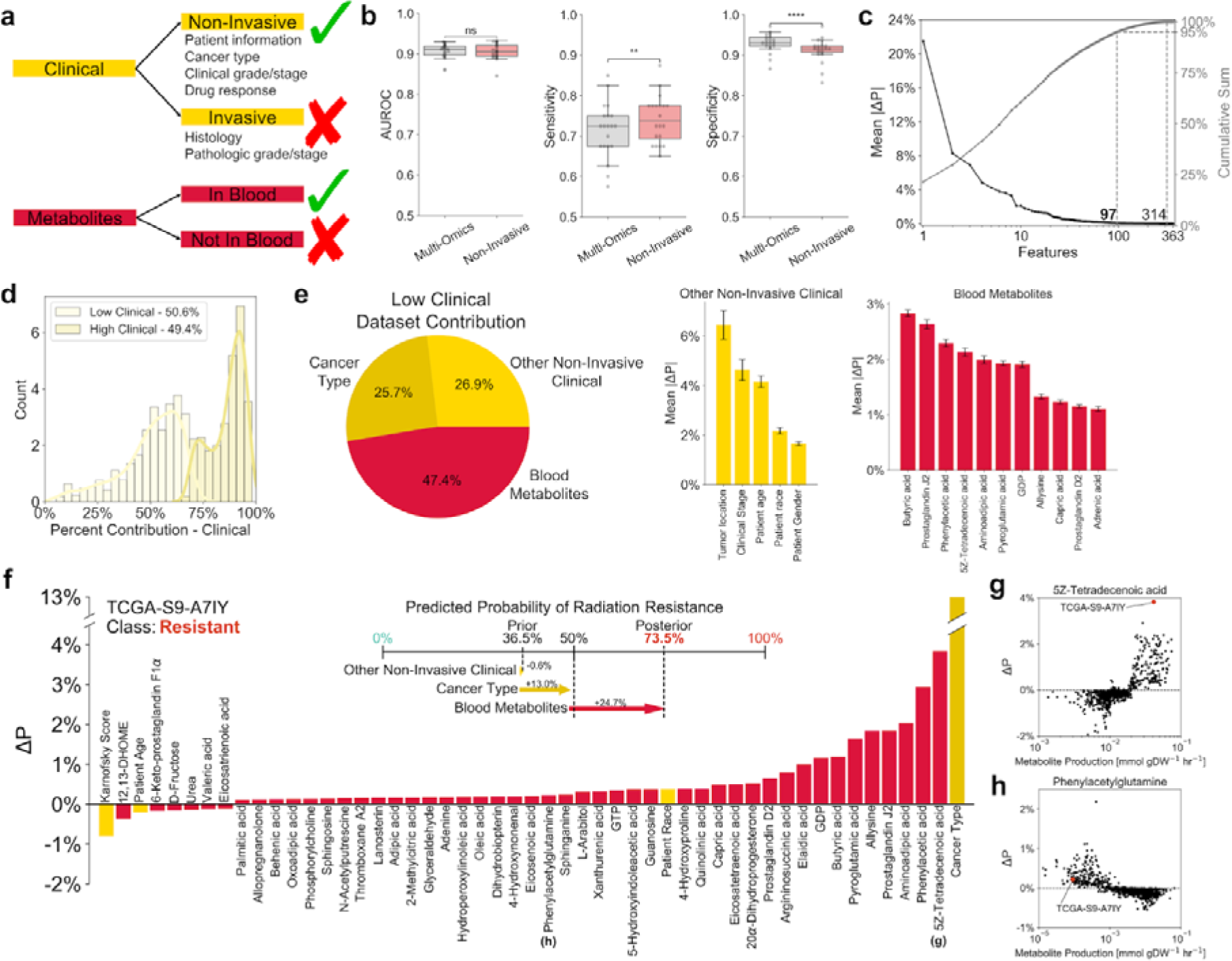
Non-invasive classifier integrating non-invasive clinical and blood-based metabolite features for prediction of radiation response. **a**, Schematic showing inclusion and exclusion criteria for features in the non-invasive classifier. **b,** Comparison of model performance between multi-omics and non-invasive classifiers. **c,** (Left, black) Feature importance scores for individual features. (Right, gray) Cumulative feature importance scores. **d,** k-means clustering of samples into “Low” and “High” clinical groups based on the relative contribution of the clinical dataset (**Supplementary Fig. 8b**). **e,** Clinical and metabolic dataset contributions among the “Low Clinical” group. Individual features with feature importance scores above 1% are shown. **f,** Breakdown of individual feature contributions towards prediction of radiation response in a representative radiation-resistant TCGA patient (TCGA-S9-A7IY). (Upper) Contribution of each dataset towards the progression from prior to posterior probability of radiation resistance. (Lower) Feature importance scores for this individual patient. **g-h,** Plots of feature importance score versus predicted metabolite production rate for two metabolic features, illustrating (**g**) a feature with large individual importance score relative to other patients (significant utility as a personalized blood-based biomarker), and (**h**) a feature with small individual importance score relative to other patients (little utility as a personalized blood-based biomarker). ns: not significant, *: p ≤ 0.05, **: p ≤ 0.01, ***: p ≤ 0.001, ****: p ≤ 0.0001.

## Discussion

Despite significant interest in methodologies for the *a priori* prediction of radiation response in cancer patients, machine learning algorithms have yet to be used in the clinical setting for informing radiation treatment^36^. Recently-developed classifiers for predicting tumor radiation response have focused mainly on gene expression data, rather than the integration of multiple -omics datasets^7, 48^. This may be in part due to a lack of metabolomics datasets from tumor biobanks including TCGA, limiting inclusion of metabolic features in machine learning classifiers for radiation response. Here, we propose a novel strategy of utilizing personalized genome-scale FBA models of radiation-sensitive and -resistant patient tumors to predict the production rates of metabolites across the Recon3D metabolic network, leveraging the accessibility of genomic and transcriptomic tumor datasets to generate metabolic insight. These metabolic features are subsequently integrated with clinical, genomic, and gene expression data from TCGA tumors to generate gene expression, multi-omics, and non-invasive classifiers for radiation response. These classifiers provide more accurate predictions of tumor radiation response compared to previously-developed classifiers, as well as novel multi-omics biomarkers associated with radiation sensitivity.

FBA model predictions of tumor metabolism and experimental validation with matched radiation-sensitive and -resistant cancer cell lines demonstrated significant re-routing of metabolic fluxes in radiation-resistant cancers, as observed by the up- and down-regulation of metabolites across multiple interconnected metabolic pathways. (**Fig. 2,5**). This flux re-routing was observed previously in the context of redox metabolism in radiation-resistant cell lines and tumors, but findings from this study suggest more widespread metabolic alterations throughout central carbon, lipid, and nucleotide metabolism^11, 49^. Our approach of systematically introducing metabolite sinks into the Recon 3D network provides a novel way of relating production fluxes to relative changes in experimentally measured metabolite levels. We observe association between increased levels of fatty acid and cholesterol metabolites with tumor radiation resistance in agreement with previous experimental evidence. Radiation-resistant head and neck cancer cells have enhanced fatty acid production from increased expression of fatty acid synthase^50^. Additionally, ionizing radiation was shown to cause increased cholesterol production in lung cancer cells^51^. Plasma levels of total and HDL cholesterol were found to be elevated in radiation-resistant SPRET/EiJ mice compared to radiation-sensitive BALB/cByJ mice, implicating cholesterol as a potential non-invasive metabolic biomarker^52, 53^. Treatment with HMG-CoA reductase inhibitors including simvastatin was reported to sensitize prostate cancer cells to radiation therapy, potentially by compromising DNA damage repair^54, 55^. Other agreements between model predictions and experimental studies include implication of inflammation-mediating eicosanoids in radiation resistance. Many prostaglandin metabolites identified in this study have previous associations with radiation resistance, and cyclooxygenase inhibitors such as aspirin may act as radiation sensitizers and improve outcomes in cervical, prostate, and rectal cancers^45, 56–58^. These findings suggest that lipid and eicosanoid metabolites may have utility as both diagnostic biomarkers as well as therapeutic targets for improving radiation response.

Integration of FBA model predictions into multi-omics machine learning classifiers for radiation response was performed by employing a dataset-independent ensemble architecture (**Fig. 3**). This approach was based on the concept of stacked generalization (having multiple “base learners” make predictions that are used as input for a separate “meta-learner”), which was shown to improve predictive accuracy in this study as well as multiple previous medical applications^59–61^. However, while in previous studies there is only one input dataset being supplied to the multiple base learners, we instead input different -omics datasets to separate base learners. The benefit of this dataset-independent approach is that the meta-learner can subsequently be used to predict which individual datasets will provide the most utility for determining radiation response in individual patients. For example, the meta-learner can accurately differentiate between “Low Clinical” patients (with large contributions of gene expression, mutation, and metabolic datasets from patient biopsy samples and genome-scale metabolic modeling) and “High Clinical” patients (with greater contribution of clinical data from electronic medical records) (**Fig. 4**). This stratification of patient populations allows for optimal resource allocation for collecting biological measurements with maximal diagnostic utility for individual cancer patients. Moreover, the use of gradient boosting machine (GBM) models as the base and meta-learners provides a significant amount of embedded feature selection; this decrease in model complexity not only lowers the cost of measuring biological features needed for prediction, but also improves the interpretability of models, increasing the likelihood of adoption by clinicians^62^.

In addition to demonstrating the utility of multi-omics data for the classification of radiation response, we found that a classifier utilizing non-invasive clinical information and blood-based metabolic biomarkers can predict radiation sensitivity with comparable accuracy (**Fig. 6**). Blood-based diagnostic tools are garnering attention for their use in early detection, monitoring, and optimal treatment identification for cancer patients^63^. Identification of novel circulating biomarkers through the integration of machine learning and genome-scale metabolic modeling could provide significant utility in adaptive radiotherapy to modify patient treatment with radiation or radiation-sensitizing chemotherapies in response to the observed efficacy of previous treatments^64^.

Although our novel approach towards integrating machine learning and genome-scale metabolic modeling for the prediction of radiation response provides many enhancements in performance accuracy and biomarker identification compared to previous studies, further improvements could yield additional benefits. Our multi-omics classifier showed that tumor histology has major impacts on both the prediction of radiation response and the clustering of “Low/High Clinical” patients, suggesting that additional features from tumor histological images could further improve classifier performance. A convolutional neural network could be added as an additional base learner with tumor H&E images from TCGA as input, providing additional histological features and identifying novel associations with radiation response^65, 66^. Additionally, changes in both DNA methylation and microRNA expression have been implicated in the tumor response to radiation therapy; inclusion of these features into the multi-omics classifier may yield additional biomarkers for improved radiation response prediction^67, 68^.

Metabolomic profiling is powerful for understanding cancer pathophysiology, identifying and monitoring clinical biomarkers, and predicting patient outcomes, but challenging to retrospectively analyze in specimen biobanks for inclusion in multi-omics data mining^69^. In this study, we demonstrate that integration of machine learning and genome-scale metabolic modeling methodologies allows for improved biomarker identification and prediction of radiation response in individual patient tumors without direct metabolomics measurements. This approach is generalizable towards other applications in guiding patient treatment, such as the prediction of chemotherapeutic response as well as identification of novel metabolic targets for pharmacological inhibition and treatment sensitization. The synergistic integration of machine learning and genome-scale metabolic modeling will inevitably yield additional insights for improving precision medicine and long-term care of cancer patients.

## Methods

### TCGA Data Retrieval and Processing

Clinical data from TCGA patients was obtained from the GDC data portal (clinical drug, clinical patient, and clinical radiation files) and the Synapse TCGA_Pancancer project (biological sample files)^2^. Drug names were standardized according to the standard available from the Gene-Drug Interactions for Survival in Cancer (GDISC) database^70^. Categorical clinical features were one-hot encoded before inputting into machine learning classifiers. RNA-Seq gene expression data was obtained from Rahman et al.’s alternative preprocessing method (GEO: GSE62944)^71^. Data from this preprocessing method showed fewer missing values, more consistent expression between replicates, and improved prediction of biological pathway activity compared to the original TCGA pipeline. Mutation data using the MuTect variant caller was obtained from the GDC data portal^2, 72^. For all data types, only features with at least two unique non-missing values were included.

### Radiation Sensitivity

TCGA samples were classified into radiation-sensitive and radiation-resistant classes according to their reported sensitivity to radiation therapy based upon the RECIST classification method. Patients with a complete or partial response to radiation (greater than 30% decrease in tumor size) were classified as radiation-sensitive, and patients with stable or progressive disease (either less than 30% decrease in tumor size, or increase in tumor size) were classified as radiation-resistant. If a patient received multiple courses of radiation therapy, they were classified based on the response to their first course.

### Data Splitting

**Supplementary Fig. 11** provides an overview of data splitting for machine learning classifier training and testing. The collection of 716 radiation-sensitive and 199 radiation-resistant samples was randomly split into training+validation (80% of all samples) and testing (20% of all samples) groups. Within the training+validation group, 5-fold cross validation was performed to optimize hyperparameter values. The training (80% of training+validation samples) group was used for training the model with a given set of hyperparameters; within this training group, 87.5% was directly used for training, and 12.5% was used to identify the iteration at which to perform early stopping during training. The validation (20% of training+validation samples) group was used to assess model performance with the given set of hyperparameters. The average validation performance across all 5 folds was used to determine the optimized set of hyperparameters; once this set was determined, the model was retrained on the entire training+validation group, and the testing group (20% of all samples) was used to assess overall model performance. 20 iterations of randomized training+validation/testing splitting were performed to analyze model predictions and performance metrics over multiple instances. All data splits were performed using stratified shuffle splitting, where the proportion of radiation-sensitive and -resistant samples was kept the same (refer to **Supplementary Fig. 11**).

### Base Learners

*N_d_* base learners were trained using an individual -omics dataset (either clinical, gene expression, mutation, or metabolic datasets), where *N_d_* is the number of individual datasets being used for the classifier. Each base learner is composed of a gradient boosting machine (GBM) model that performs two-class classification (predicting either radiation sensitivity or resistance for each patient) using features from an individual dataset, such as clinical, genomics, transcriptomics. GBM models using decision tree ensembles have many useful characteristics compared to other machine learning algorithms, including embedded feature selection, capability of handling missing values (which is common in clinical datasets), and efficient management of high-dimensional datasets (where the number of features greatly exceeds the number of samples)^73, 74^. XGBoost (v0.90) was used to develop GBM base learners and meta-learners^75^.

Bayesian optimization was performed to optimize hyperparameter values for each GBM model. At each iteration of Bayesian optimization, 5-fold cross validation was used to calculate the performance of a particular set of hyperparameters. Weighted log loss was used as the performance metric for both GBM model training and evaluating model performance on validation sets:

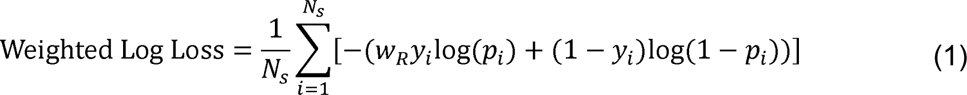

where *y_i_* is the true class label of sample *i* (*y_i_*=0 if sensitive, *y_i_*=1 if resistant), *p_i_* is the predicted probability of sample *i* being radiation-resistant (belonging to class 1), *w_R_* is the weight given to radiation-resistant samples (*w_R_* = # sensitive samples / # resistant samples), and *N_S_* is the total number of samples. The weight given to radiation-resistant samples accommodates for the fact that there are more radiation-sensitive samples than radiation-resistant samples, and prevents classifiers from focusing on optimizing performance exclusively on radiation-sensitive samples. The mean weighted log loss plus one standard error over all 5 folds of cross-validation is used to choose the hyperparameter set with best performance. During model training, early stopping is employed to prevent overfitting.

For individual samples, each of the *N_d_* base learners outputs the predicted probability of radiation resistance (*p_1_*, *p_2_*, …, *p_Nd_*) using features from the individual data type. Each base classifier receives the same training/validation/testing split of samples.

### Meta-learner

For every sample within the 5 validation sets used for the base learners, each base learner’s output prediction of radiation resistance (*p_i_*) is compared to the sample’s true radiation response class (*y_i_*). The meta-learner is trained to predict the optimal base learner which provides the most accurate prediction of radiation response for each sample, based on the sample’s multi-omics features. This meta-learner performs an *N_d_*-class classification, where *N_d_* is the number of independent base learners. The features this meta-learner is trained on include all features from the *N_d_* datasets which have non-zero feature importance scores from their respective base learners; features which do not impact base learner predictions are not included, which increases the training speed while maintaining meta-learner accuracy. Because validation samples from the 5-fold cross validation were not directly used in base learner training, they can be used to train this meta-learner without overfitting or inflation of model performance metrics.

Implementation of the meta-learner is analogous to that of each base learner, using a GBM model, Bayesian optimization, early stopping, and 5-fold cross validation. Multiclass log loss was used as the performance metric for both GBM model training and evaluating model performance ^76^:

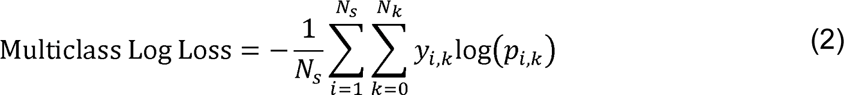

where *y_i,k_* is 1 if dataset *k* is the true optimal dataset of sample *i* and 0 otherwise, *p_i,k_* is the predicted probability of dataset *k* being the optimal dataset of sample *i*, *N_S_* is the total number of samples, and *N_k_* is the total number of datasets. The mean multiclass log loss plus one standard error over all 5 folds of cross-validation is used to choose the optimal hyperparameter set with best performance.

For individual samples, the meta-learner outputs *N_d_* probabilities (*w_1_*, *w_2_*, …, *w_Nd_*) that each base learner is optimal for that sample (all *N_d_* probabilities sum to 1). Note that, once the meta-learner is trained using the predicted probabilities from the base learners, the base learners and meta-learner act independently of each other when used on new testing samples.

### Radiation Response Prediction

Each testing sample is run through 1) all *N_d_* base learners to obtain the predicted probabilities of radiation resistance using each of the *N_d_* individual datasets (*p_1_*, *p_2_*, …, *p_Nd_*), and 2) the meta-learner to obtain the predicted probabilities that each of the *N_d_* base learners/datasets is optimal for that sample (*w_1_*, *w_2_*, …, *w_Nd_*). To obtain the final predicted probability of radiation resistance for the testing sample, the weighted average of the base learner probabilities is taken, with the meta-learner probabilities as weights:

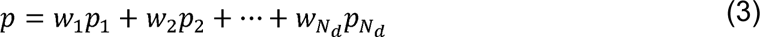

Samples with *p* < 0.5 are classified as radiation-sensitive, while samples with p > 0.5 are classified as radiation-resistant.

### Bayesian Optimization

Bayesian optimization was used to optimize GBM hyperparameters for both the base learner and meta-learner classifiers. This iterative approach automates the search for hyperparameter values by calculating an acquisition function which provides the expected benefit of sampling a particular point in hyperparameter space on the overall search for hyperparameters with minimal cross-validation error. At each iteration, the point in hyperparameter space with the largest acquisition function value is chosen, 5-fold cross validation is used to determine the performance of those particular hyperparameters, and the acquisition function is updated to then determine which next point in hyperparameter space will be sampled. Hyperopt (v0.1.2) was used to perform Bayesian optimization^77^. **Supplementary Table 2** provides the 8 GBM hyperparameters chosen for optimization of both base learner and meta-learner classifiers, with the ranges of values in the hyperparameter search space. 2^8^=256 iterations of Bayesian optimization were performed for each classifier.

### Classifier Performance Metrics

Final classifier performance was assessed on testing samples across the 20 iterations of randomized training+validation/testing splitting. The following performance metrics were used:

1. Weighted log loss: equation (1)
2. Area under the receiver operating characteristic curve (AUROC)
3. Balanced accuracy, an accuracy metric that corrects for unequal numbers of radiation-sensitive and -resistant patients:

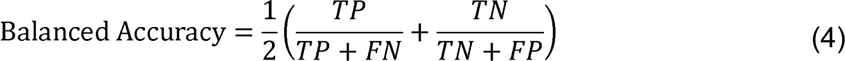
4. Sensitivity:

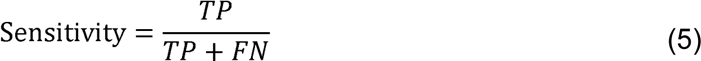
5. Specificity:

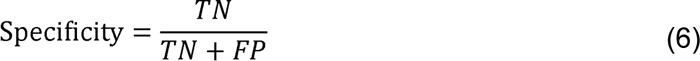
6. Positive predictive value:

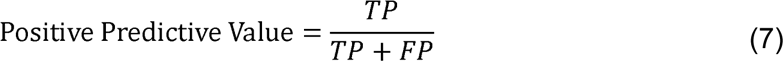
7. Negative predictive value:

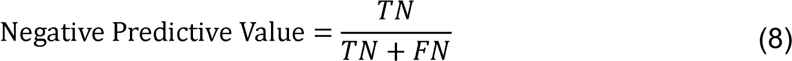

### Feature Importance Scores

The importance of individual features towards the prediction of radiation response, both averaged across all samples as well as for individual samples, was determined by calculating Shapley Additive Explanations (SHAP) values for each classifier. Each SHAP value represents the change in predicted probability of radiation resistance for patient *i* attributed to feature *j*^78^. Features with positive SHAP values for patient *i* signify those where the particular value of feature *j* attributed to patient *i* is such that it increases patient *i*’s predicted probability of radiation resistance. Larger absolute SHAP values indicate features with larger overall contributions (either negatively or positively). Mean absolute SHAP values across all samples provide an indication of the overall importance of a particular feature in the classifier’s prediction of radiation response. SHAP values were averaged across 20 training+validation/testing splits by a weighted average ^79^, with weights proportional to the inverse of the weighted log loss performance metric on the testing set for that split. This weighted average allows model analysis to be more reflective of the more accurate predictions, so that identified biomarkers are more likely to be true biomarkers rather than artifacts of poorly-performing predictions. Values were normalized by the difference between prior and posterior probabilities of radiation resistance for each sample. SHAP v0.29.1 was used to calculate SHAP values^80^.

### Comparison of Machine Learning Algorithms

scikit-learn v0.21.2 functions sklearn.ensemble.RandomForestClassifier() and sklearn.linear_model.LogisticRegression() were used to implement random forest and logistic regression with L1 regularization classifiers, respectively^81^. Keras v2.3.1 was used to implement the neural network with L1 regularization classifier (https://github.com/keras-team/keras). Weighted log loss (equation (1)) was used as the loss function for the neural network classifier, and early stopping was performed. Missing values were imputed and scaled using sklearn.impute.SimpleImputer() and sklearn.preprocessing.StandardScaler() functions, respectively, before training with the random forest, logistic regression, and neural network classifiers. **Supplementary Tables 3-5** provide the hyperparameters and value ranges used for Bayesian optimization with each algorithm; 256 iterations of Bayesian optimization were performed for each classifier. Each classifier, including the GBM classifier, was run using the same training, validation, and testing samples at each of 20 training+validation/testing splits so that performance can be accurately compared.

### Comparison of Gene Expression Datasets

11 gene expression sets for oxic radiation response gene expression sets in the RadiationGeneSigDB database were compared to the set of 782 significant genes from the gene expression classifier^7^. Gene names from RadiationGeneSigDB gene sets were converted to Entrez gene ID’s and gene symbols. Those genes where a matching Entrez gene ID or gene symbol could not be found were removed. Additionally, those genes that were not in both TCGA and CCLE gene expression datasets were removed.

To compare performance of gene expression sets on TCGA data, classification models were trained to predict radiation-sensitive or -resistant classes of TCGA tumor samples using gene expression data from only the subset of genes for an individual set. Model performance was assessed using weighted log loss (equation (1)) and AUROC metrics. To compare performance of gene expression sets on CCLE data, regression models were trained to predicted radiation response (reported as area under the curve of survival vs. radiation dose) of CCLE cell lines using gene expression data from only the subset of genes for an individual set^82^.Model performance was assessed using mean absolute error (MAE) and mean squared error (MSE) metrics.

### Flux Balance Analysis (FBA)

Generation of personalized FBA models of individual TCGA tumor samples was performed as described in Lewis et al^11^. To predict the maximum production of a particular metabolite in FBA models, the following objective function was used:

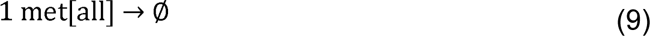

where “met” is the metabolite to be maximized, and “[all]” represents the maximization of this objective function across all cellular compartments where the metabolite is located. This creates an artificial sink for a particular metabolite in the Recon3D metabolic network, resulting in the maximization of reaction fluxes generating this metabolite. We hypothesized that this objective function would be valid and thus yield accurate predictions for metabolites with large differences in production between radiation-sensitive and -resistant tumors, as these would be particularly beneficial to either tumor class and thus the metabolic network of these tumors would be optimized to maximize levels of the metabolite.

The modeled external compartment contained all metabolites found in DMEM/F-12 cell culture media (Thermo Fisher Scientific, Cat#11320) as well as fetal bovine serum (FBS) to match the cell culture media used for experimental validation^83^. All 871 metabolites in the Recon3D human metabolic reconstruction that (1) had KEGG database ID’s, (2) were not present in the extracellular media, and (3) were capable of being produced by all FBA tumor models, were included in the FBA metabolite production screen.

### NCI-60 Data Retrieval and Processing

Experimental metabolomics data from NCI-60 cancer cell lines was obtained from the Developmental Therapeutics Program (DTP) of the National Cancer Institute (NCI) (https://wiki.nci.nih.gov/display/NCIDTPdata/Molecular+Target+Data). Normalized concentration entries without metabolite names or for isobars were excluded. Cell line surviving fraction at 2 Gy radiation (SF2) values were obtained from Amundson et al ^32^.

### Cell Culture

**Supplementary Table 1** provides the matched radiation-sensitive and radiation-resistant cell lines used for experimental validation of metabolite levels predicted from FBA models. All cell lines were maintained in DMEM/F-12 cell culture media (Thermo Fisher Scientific, Cat#11320) with 10% FBS (Sigma-Aldrich, Cat#F4135) at 37°C and 5% CO_2_, and were free of *Mycoplasma*.

### Metabolomics

Three biological replicates of each cell line were grown in separate T-25 flasks with the cell culture conditions described above. Cell pellets with approximately 1 million cells were obtained from trypsinization, centrifugation, and removal of supernatant. Samples were reconstituted in 90% MeOH, 10% H_2_O at a ratio of 200 μL/1 million cells. Aliquots of the supernatant were combined to create a pooled sample used for quality control. Aliquots of the samples were transferred to LC vials and stored at 4°C.

Hydrophilic interaction chromatography-tandem mass spectrometry (HILIC-MS/MS) untargeted metabolomics was performed. Chromatography parameters were as follows: BEH HILIC Column, 150 mm X 2.1 mm, 1.7 μm; mobile phase A: 80% H2O / 20% ACN, 10mM ammonium formate, 0.1% FA; mobile phase B: 100% ACN, 0.1% FA; column temperature: 40°C; 2 μ sample injection. MS parameters were as follows: resolution: 240,000; scan range: 70-1050 m/z; polarity: positive/negative; AGC target: 1e5. MS^2^ parameters were as follows: isolation window: 0.8 m/z; detector: Orbitrap; polarity: positive/negative; fragmentation method: HCD; collision energy: 15, 30, 45; resolution: 30,000.

Compound Discoverer 3.1 was used to perform quality control, putative metabolite identification, and quantification of metabolite levels. Results for positive and negative ion modes were combined. Metabolites with no identified name were removed from the analysis. If duplicate metabolites with the same identification were obtained, then the entry with the largest maximum area was used. KEGG ID’s for each metabolite were manually identified based on metabolite name, molar mass, and chemical formula. Metabolites from experimental metabolomics were matched to those from FBA analysis by matching KEGG ID’s.

For the comparison of model-predicted and experimentally-measured metabolite levels, all metabolites within the following Recon3D subsystems that were matched with experimental metabolites were included in the analysis:

● Nucleotide Metabolism: “Nucleotide interconversion”, “Nucleotide salvage pathway”, “Pentose phosphate pathway”, “Purine catabolism”, “Purine synthesis”, “Pyrimidine catabolism”, “Pyrimidine synthesis”
● Lipid Metabolism: “Cholesterol metabolism”, “Fatty acid oxidation”, “Fatty acid synthesis”, “Glycosphingolipid metabolism”, “Phosphatidylinositol phosphate metabolism”, “Sphingolipid metabolism”, “Steroid metabolism”
● Cysteine/Antioxidant Metabolism: “Glutathione metabolism”, “ROS
detoxification”, plus metabolite “Lipoamide”
● Immune System Mediators: “Arachidonic acid metabolism”, “Eicosanoid metabolism”

### Code and Data Availability

Jupyter notebooks containing Python code for running and analyzing the gene expression, multi-omics, and non-invasive classifiers for radiation response are available at https://github.com/kemplab/ML-radiation. Additionally, the gene sets and code used to compare the significant gene list from our gene expression classifier to those from the RadiationGeneSigDB database are available.

The following datasets are available at https://github.com/kemplab/ML-radiation:

- *Dataset 1.* Feature importance scores (ΔP) from the gene expression classifier, for individual TCGA patients
- *Dataset 2.* FBA model-predicted metabolite production rates in TCGA tumors
- *Dataset 3.* Experimental metabolomics data from radiation-sensitive and - resistant cancer cell lines
- *Dataset 4.* Comparison of model-predicted and experimentally-validated metabolite levels in radiation-sensitive and -resistant cancers
- *Dataset 5.* Feature importance scores (ΔP) from the multi-omics classifier, for individual TCGA patients
- *Dataset 6.* Feature importance scores (Δ from the non-invasive classifier, for individual TCGA patients

## Acknowledgements

The authors gratefully acknowledge support for this work from an NIH/NCI F30 CA224968 fellowship (PI: J.E.L.; Sponsor: M.L.K.) and an NIH/NCI U01 CA215848 grant (PI: M.L.K.). We wish to acknowledge David Gaul and the core facilities at the Parker H. Petit Institute for Bioengineering and Bioscience at the Georgia Institute of Technology for the use of their shared equipment, services and expertise.

## Author Contributions

Conceptualization, J.E.L. and M.L.K.; Methodology, J.E.L.; Software, J.E.L.; Validation, J.E.L.; Formal Analysis, J.E.L.; Investigation, J.E.L.; Resources, M.L.K.; Data Curation, J.E.L.; Writing – Original Draft, J.E.L.; Writing – Review & Editing, J.E.L. and M.L.K.; Visualization, J.E.L.; Supervision, M.L.K.; Project Administration, J.E.L. and M.L.K.; Funding Acquisition, J.E.L. and M.L.K.

## Supplementary Figures

**Supplementary Fig. 1.**
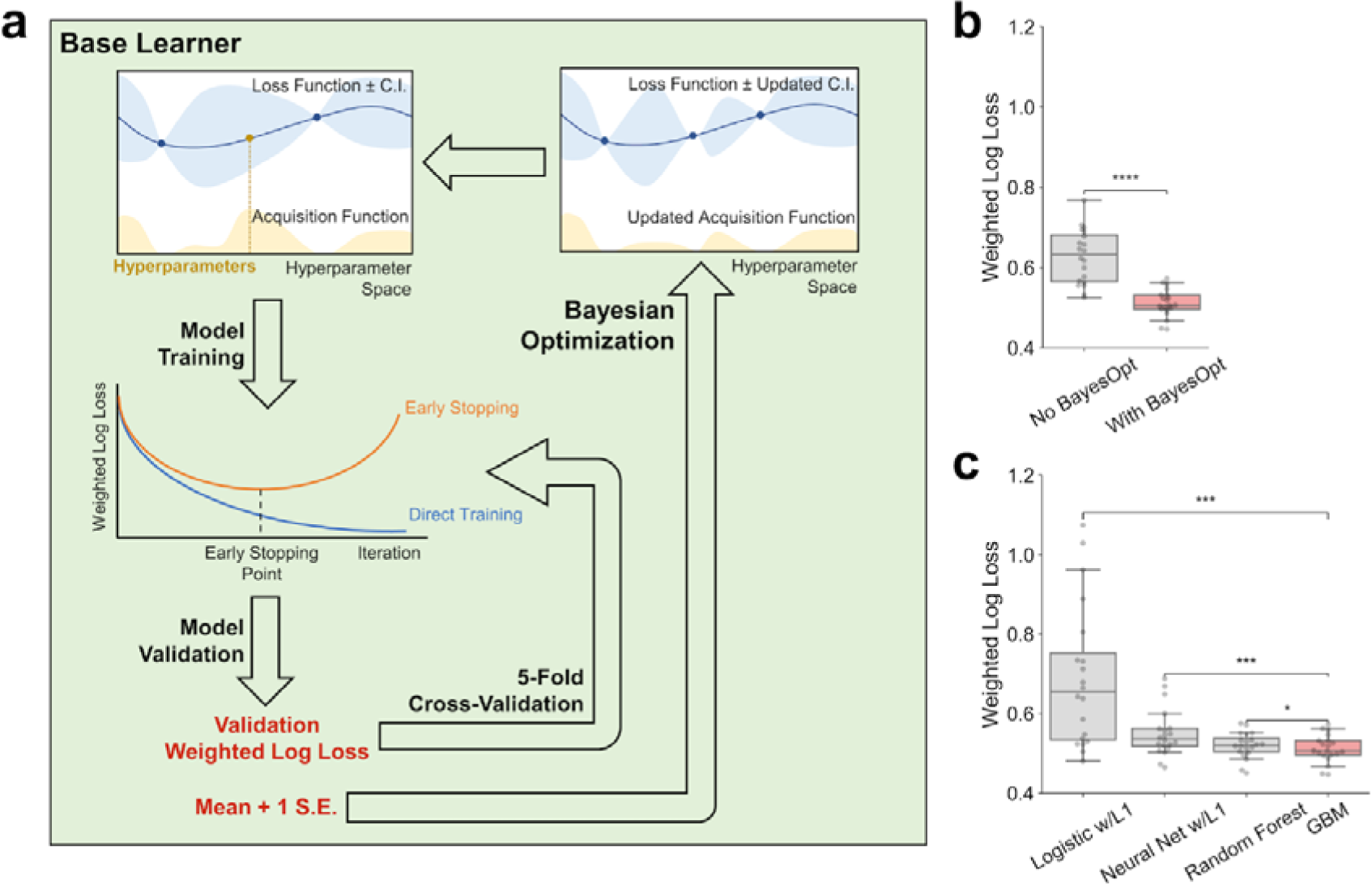
Base learner used in the gene expression, multi-omics, and non-invasive classifiers for radiation response. **a**, Base learner performing two-class classification of radiation response, utilizing a gradient boosting machine (GBM) algorithm with Bayesian optimization, early stopping, and 5-fold cross validation to determine optimal hyperparameter values. Note that for the gene expression classifier, this base learner is not integrated with other base learners or a meta-learner as only one dataset is utilized. **b-c,** Performance of classifier trained on gene expression data from TCGA tumors, comparing (**b**) use of Bayesian optimization compared to no Bayesian optimization, and (**c**) GBM-based classifiers versus other machine learning algorithms. ns: not significant, *: p ≤ 0.05, **: p≤ 0.01, ***: p≤0.001, ****: p≤0.0001.

**Supplementary Fig. 2.**
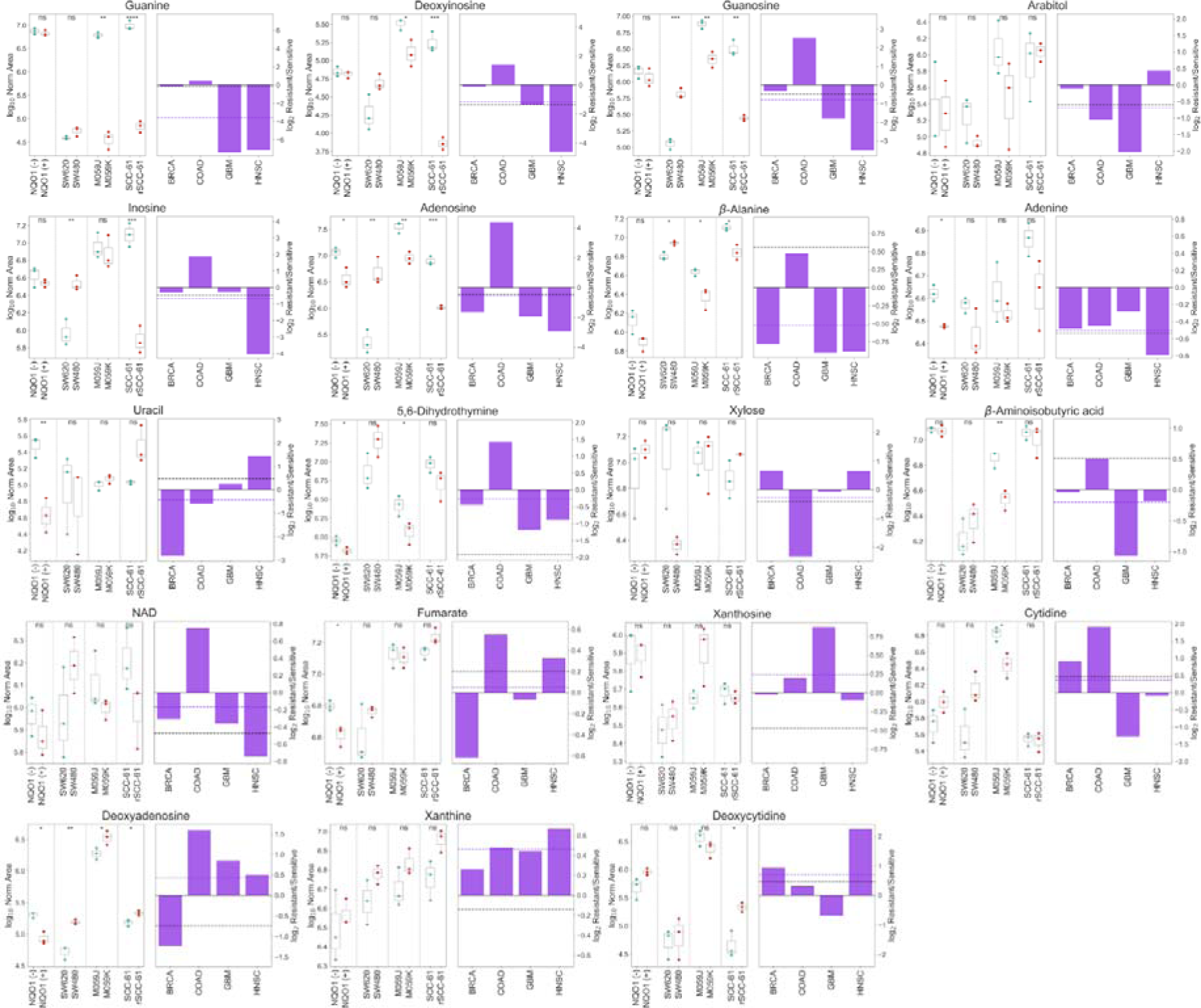
Experimentally-measured concentrations of nucleotide metabolites in matched radiation-sensitive and -resistant cell lines. (Left) Replicate metabolite concentrations from all four cell line pairs, with values expressed as the log_10_ normalized area from LC-MS/MS. ns: not significant, *: p ≤ 0.05, **: p≤ 0.01, ***: p 0.001, ****: p ≤ 0.0001. (Right) Bars: Ratio value for each≤ cell line pair, expressed as the log_2_ ratio of mean radiation-resistant concentration versus mean radiation-sensitive concentration. Colored line: mean experimental log_2_ Resistant/Sensitive across all four cell line pairs. Black line: FBA model-predicted log_2_ ratio of average metabolite production in radiation-resistant versus -sensitive TCGA tumors.

**Supplementary Fig. 3.**
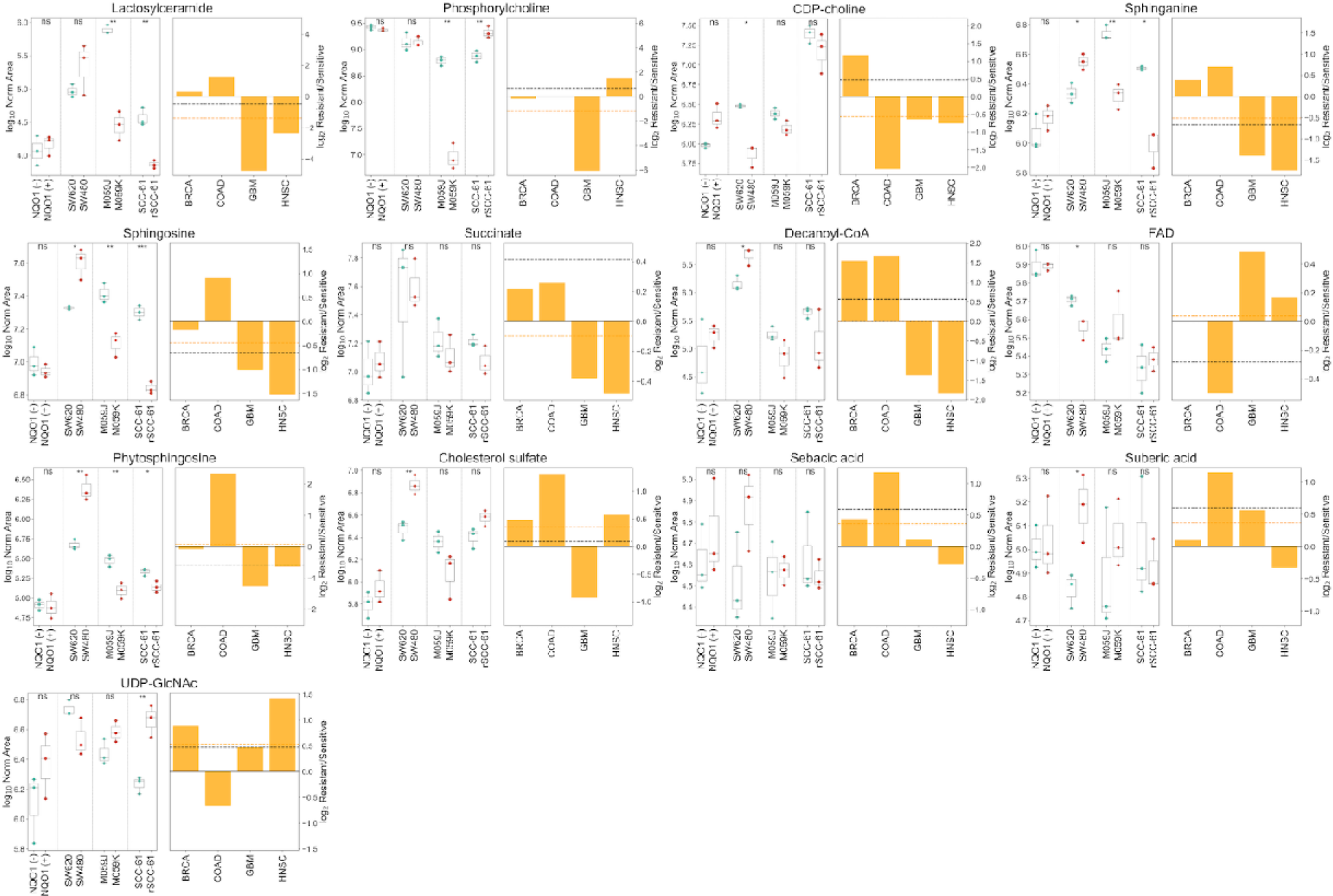
Experimentally-measured concentrations of lipid metabolites in matched radiation-sensitive and -resistant cell lines. (Left) Replicate metabolite concentrations from all four cell line pairs, with values expressed as the log_10_ normalized area from LC-MS/MS. ns: not significant, *: p ≤ 0.05, **: p≤ 0.01, ***: p 0.001, ****: p ≤ 0.0001. (Right) Bars: Ratio value for each≤ cell line pair, expressed as the log_2_ ratio of mean radiation-resistant concentration versus mean radiation-sensitive concentration. Colored line: mean experimental log_2_ Resistant/Sensitive across all four cell line pairs. Black line: FBA model-predicted log_2_ ratio of average metabolite production in radiation-resistant versus -sensitive TCGA tumors.

**Supplementary Fig. 4.**
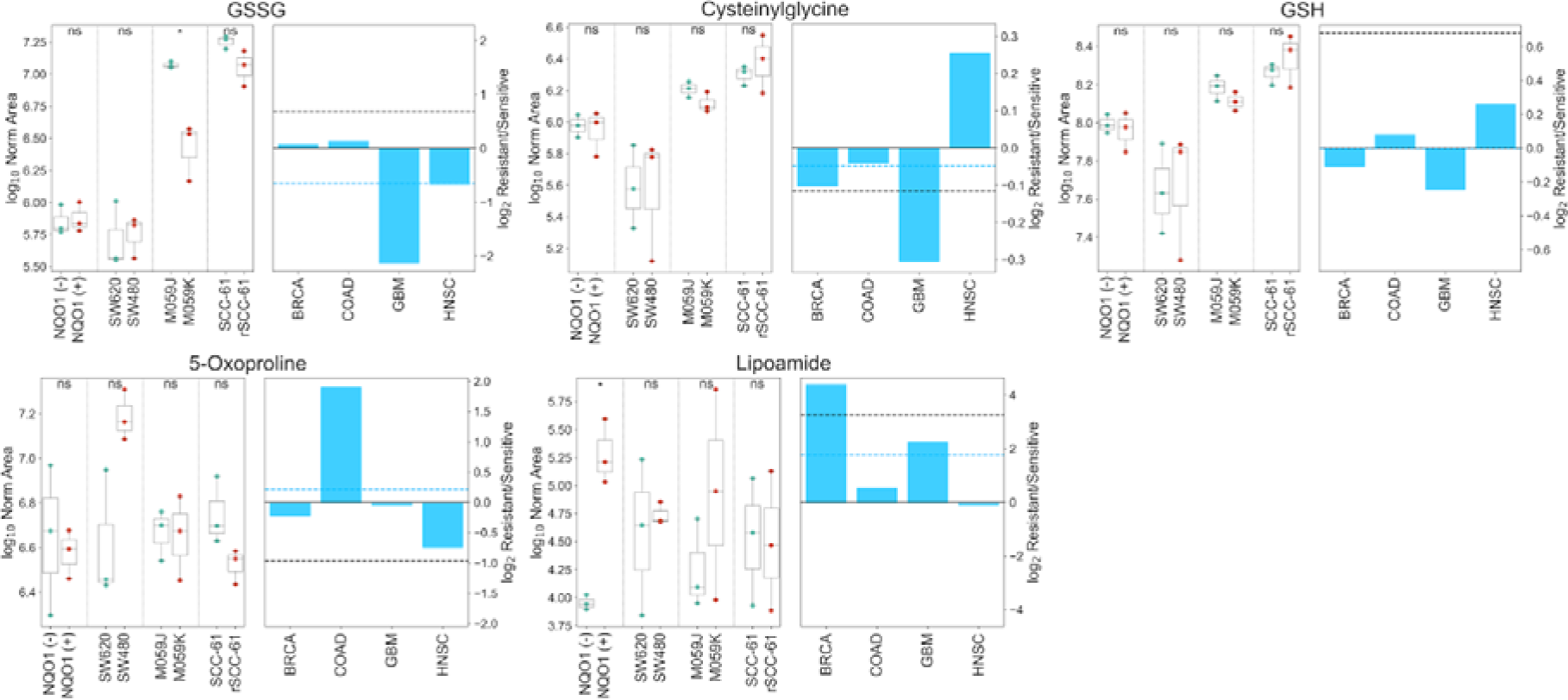
Experimentally-measured concentrations of cysteine/antioxidant metabolites in matched radiation-sensitive and -resistant cell lines. (Left) Replicate metabolite concentrations from all four cell line pairs, with values expressed as the log10 normalized area from LC-MS/MS. ns: not significant, *:p 0.05, **: p ≤ ≤ 0.01, ***: p≤ 0.001, ****: p≤ 0.0001. (Right) Bars: Ratio value for each cell line pair, expressed as the log_2_ ratio of mean radiation-resistant concentration versus mean radiation-sensitive concentration. Colored line: mean experimental log_2_ Resistant/Sensitive across all four cell line pairs. Black line: FBA model-predicted log_2_ ratio of average metabolite production in radiation-resistant versus -sensitive TCGA tumors.

**Supplementary Fig. 5.**
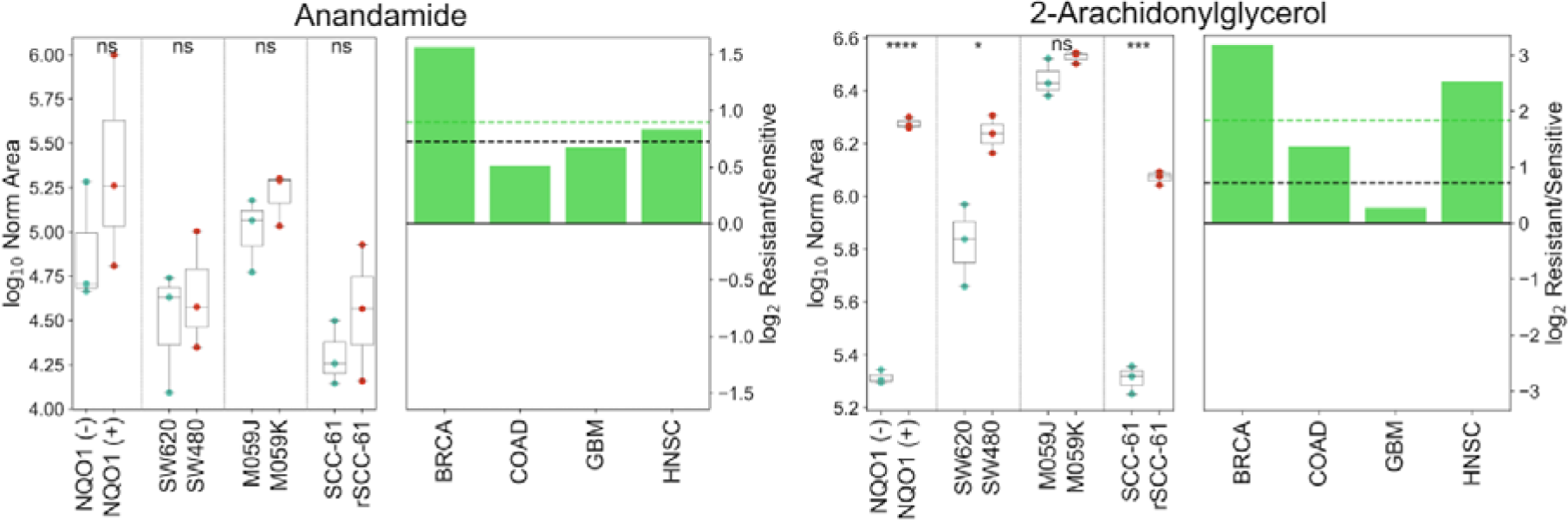
Experimentally-measured concentrations of immune system mediating metabolites in matched radiation-sensitive and -resistant cell lines. (Left) Replicate metabolite concentrations from all four cell line pairs, with values expressed as the log10 normalized area from LC-MS/MS. ns: not significant, *: p 0.05, **: p ≤ ≤ 0.01, ***: p≤ 0.001, ****: p≤ 0.0001. (Right) Bars: Ratio value for each cell line pair, expressed as the log_2_ ratio of mean radiation-resistant concentration versus mean radiation-sensitive concentration. Colored line: mean experimental log_2_ Resistant/Sensitive across all four cell line pairs. Black line: FBA model-predicted log_2_ ratio of average metabolite production in radiation-resistant versus -sensitive TCGA tumors.

**Supplementary Fig. 6.**
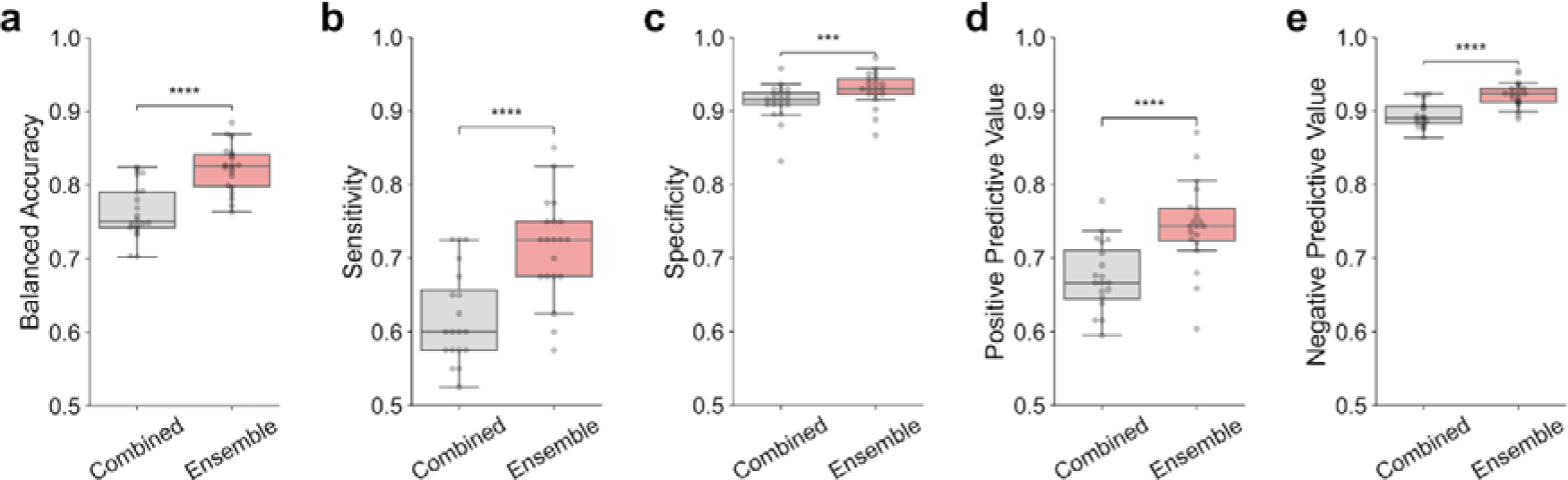
Performance of the multi-omics classifier, comparing the dataset-independent ensemble architecture versus combining datasets together before training on a single classifier. Multiple alternative classifier performance metrics are provided.

**Supplementary Fig. 7.**
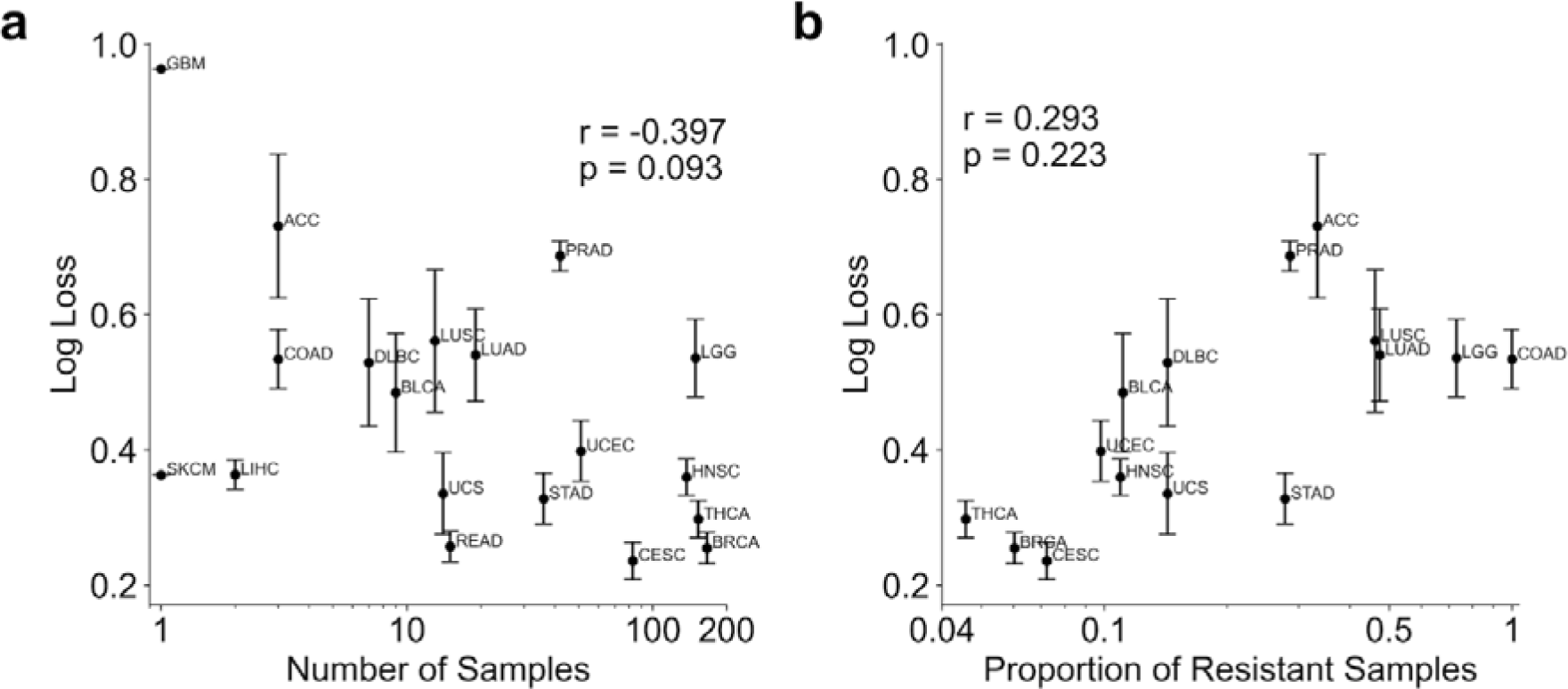
Comparison of multi-omics classifier performance on samples from different cancer types. a, Correlation between sample log loss and number of samples within each cancer type. **b,** Correlation between sample log loss and proportion of radiation-resistant samples within each cancer type.

**Supplementary Fig. 8.**
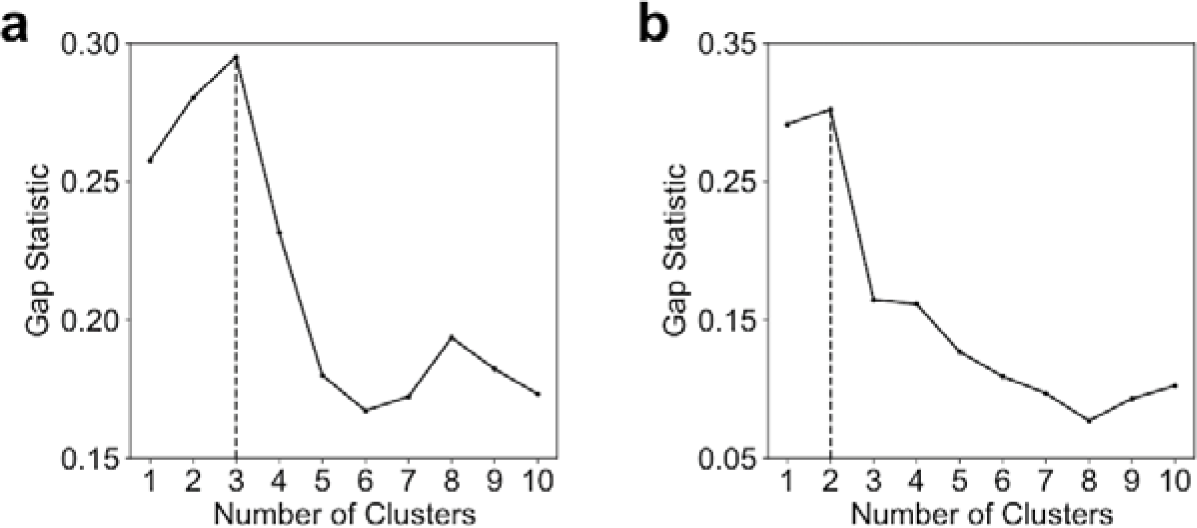
*k*-Means clustering of clinical dataset contributions for individual samples. Gap statistic values for each value of *k* are shown for the (**a**) multi-omics classifier, and (**b**) non-invasive classifier.

**Supplementary Fig. 9.**
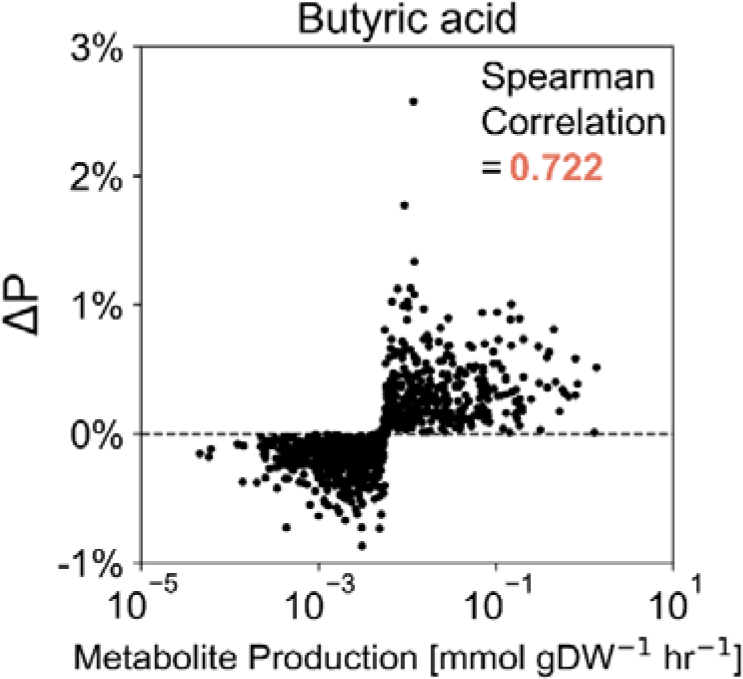
Regression between feature importance score and predicted metabolite production rate for a representative metabolite. Values are shown for each individual patient tumor.

**Supplementary Fig. 10.**
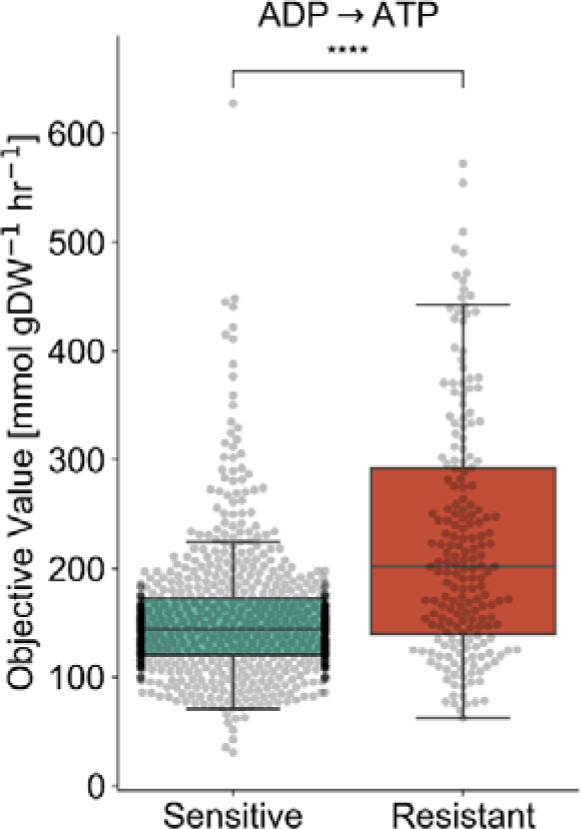
Comparison of FBA model-predicted maximal conversion of ADP to ATP between radiation-sensitive and -resistant. TCGA tumors. ****: p≤0.0001.

**Supplementary Fig. 11.**
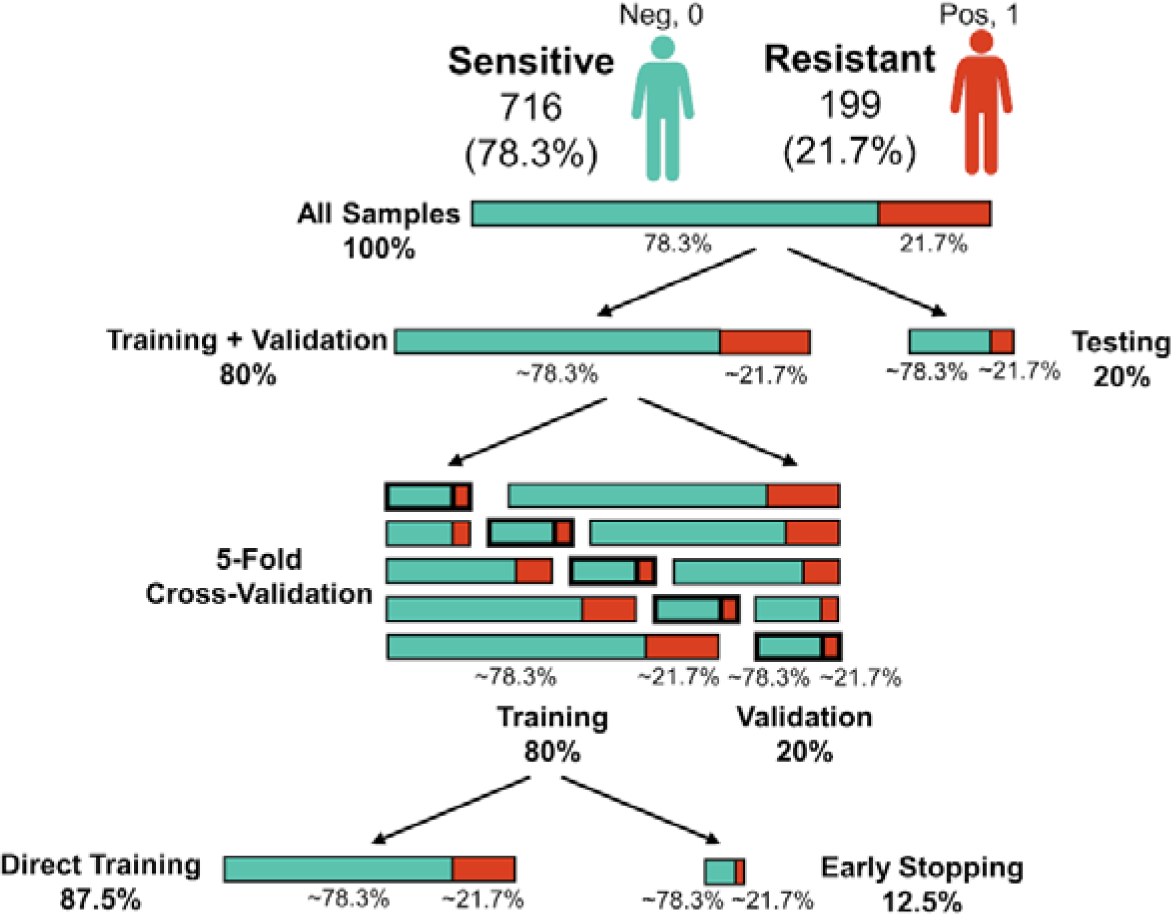
**Data splitting for classifier training and testing.**

## Supplementary Tables

**Supplementary Table 1.**
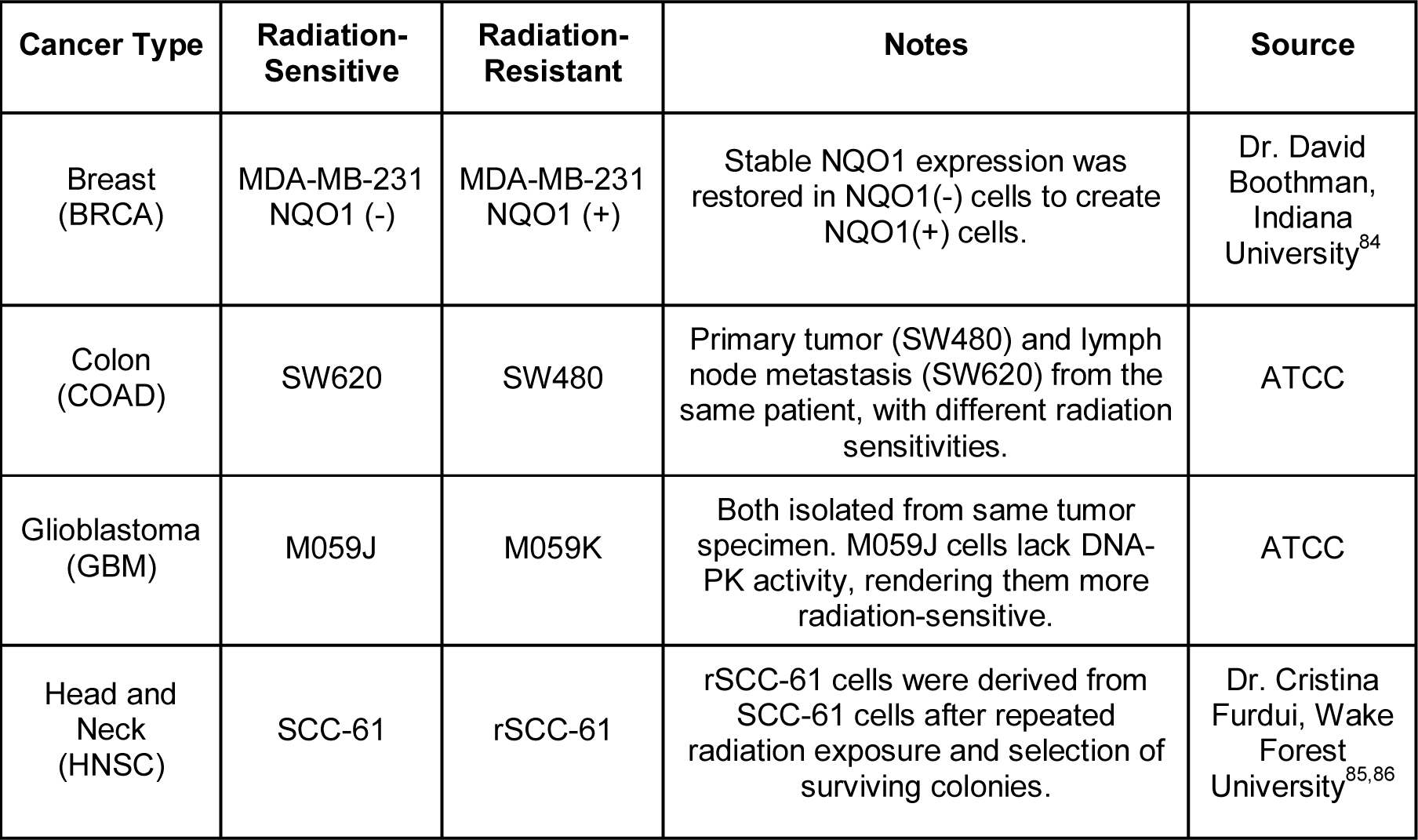
Matched radiation-sensitive and radiation-resistant cancer cell lines

**Supplementary Table 2.** Hyperparameter ranges for Bayesian optimization with gradient boosting machine classifiers.

| Parameter Name | Distribution | Lower Bound | Upper Bound |
| --- | --- | --- | --- |
| eta | log-uniform | 0.01 | 0.5 |
| gamma | log-uniform | 0 | 5 |
| max_depth | uniform | 1 | 11 |
| subsample | uniform | 0.5 | 1 |
| colsample_bytree | uniform | 0.5 | 1 |
| colsample_bylevel | uniform | 0.5 | 1 |
| reg_lambda | log-uniform | 1 | 4 |
| reg_alpha | log-uniform | 0 | 1 |

**Supplementary Table 3.** Hyperparameter ranges for Bayesian optimization with the random forest classifier.

| Parameter Name | Distribution | Lower Bound | Upper Bound |
| --- | --- | --- | --- |
| n_estimators | log-uniform | 1 | 1000 |
| criterion | choice | entropy, gini |  |
| max_depth | uniform | 1 | 11 |
| min_samples_split | uniform | 0 | 1 |
| min_samples_leaf | uniform | 0 | 0.5 |
| max_features | choice | log2, sqrt |  |

**Supplementary Table 4.** Hyperparameter ranges for Bayesian optimization with the logistic regression classifier with L1 regularization.

| Parameter Name | Distribution | Lower Bound | Upper Bound |
| --- | --- | --- | --- |
| C | log-uniform | 0.001 | 1000 |

**Supplementary Table 5.** Hyperparameter ranges for Bayesian optimization with the neural network classifier with L1 regularization.

| Parameter Name | Distribution | Lower Bound | Upper Bound |
| --- | --- | --- | --- |
| Number of layers | uniform | 1 | 5 |
| Neurons per layer | choice | 8, 16, 32, 64, 128, 256, 512, 1024 |  |
| activation_function | choice | elu, relu, sigmoid |  |
| optimizer | choice | adam, rmsprop, sgd |  |
| l1 | log-uniform | 0.000001 | 0.1 |
| dropout | uniform | 0 | 0.5 |

